# Sfh3 is a sterol-modulated PI/PI4P transfer protein that confers azole resistance in yeast by specifically mediating lanosterol/PI exchange

**DOI:** 10.64898/2026.10.26.761302

**Authors:** Amazigh Hamaï, Pauline Perné, Abhishek Tripathi, Camille Syska, Anna Cardinal, Vanessa Delfosse, Alenka Čopič, Natarajan Arul Murugan, Delphine Debayle, Guillaume Drin

## Abstract

Lipid transfer proteins precisely distribute lipids within eukaryotic cells, supporting functions such as vesicular trafficking and cellular signaling. Here, we analyzed whether Sfh3 (Sec14 homolog 3) functions as a sterol/phosphatidylinositol (PI) exchanger to clarify its role in neutral lipid mobilization from lipid droplets and in azole resistance in yeast. Using real-time assays, we found that its ergosterol transfer capacity is not activated by PI and is inhibited by ergosterol biosynthesis intermediates, such as lanosterol. Next, we found that Sfh3 exhibits both PI and PI4P transfer activities, which sterols can modulate. Through kinetic, structure-function, and thermal stability analyses, we identified remarkable Sfh3 mutants and ranked Sfh3’s relative affinity for its lipid ligands. This allowed us to show that Sfh3 performs selective lanosterol/PI exchange, provided it recognizes PI, in the presence of ergosterol, and that this activity is necessary for yeast resistance to miconazole. Collectively, our data show that Sfh3 is a sterol-regulated PI/PI4P transfer protein, potentially linking sterol metabolism and PI4P signaling during lipid droplet utilization. In azole-treated yeast, elevated lanosterol levels may shift Sfh3 into an exclusive PI/lanosterol exchange mode, promoting cellular detoxification and conferring drug resistance. By revealing unanticipated Sfh-mediated lipid transfer mechanisms, our study provides new insights into the Sec14 family.

## Introduction

In eukaryotic cells, lipids are highly diverse and precisely distributed between organelle membranes and the plasma membrane (PM)^1^, as well as across the two leaflets of these membranes, often asymmetrically^2^. This distribution is crucial for most cellular functions, such as vesicular trafficking and cell signaling^3–5^. It is now well established that intracellular lipid distribution relies on cytosolic proteins, known as lipid transfer proteins (LTPs)^6^, that move lipids between membranes. Some of these proteins, belonging to the Sec14 family, the phosphatidylinositol (PI)-transfer protein (PITP) family, or the Oxysterol-binding protein-related proteins (ORPs) family, critically impact cellular functions by exchanging two different lipid ligands between membranes^7–8^.

Sec14, the prototypical member of the eponymous family, is a ∼35 kDa yeast protein containing a lipid-binding cavity that can harbor either phosphatidylcholine (PC) or PI. Pioneering works established that Sec14 mediates PC/PI exchange to support PI4P biosynthesis, a key phosphoinositide in Golgi signaling, required for post-Golgi vesicle trafficking and secretory competence in yeast^7–8^. After nearly four decades since its discovery, it remains unclear whether Sec14 transfers PI from its site of synthesis, the endoplasmic reticulum (ER), to the Golgi to maintain a local PI pool that is then converted by Golgi-resident PI 4-kinases into PI4P, or whether it directly performs PI/PC exchange on the Golgi membrane to present PI to PI 4-kinases to boost their catalytic activity^7–8^.

Compared with Sec14, even less is known about many proteins in Eukaryotia that contain a Sec14 domain (also called CRAL-TRIO) capable of binding lipids as well as non-lipid ligands^9–10^. Human cells encode almost 30 different proteins, often with multidomain organization, including a Sec14 domain, and some are linked to diseases such as cancer, blindness, and neurodegeneration. Yeast cells express five Sec14 homologs, called Sfh proteins, which bind PI and a second ligand that is not necessarily PC. Computational analyses of this small subset of proteins indicated that variations in the Sec14 domain define specific cavity microenvironments to host different ligands, while conservation of a few residues at specific positions allows the recognition of PI^11^. How Sfh proteins and, more generally, Sec14-domain-containing proteins translate their capacity to recognize different ligands into cellular functions remains an active field of research^8, 12^

To better understand the biology of Sec14-domain-containing proteins, we focused on Sfh3 (a.k.a. Pdr16), which has received a lot of attention recently as it is associated with processes that regulate energy utilization from lipid droplets (LDs) in yeast ^13–15^ and was otherwise long known to confer drug resistance in yeast ^16–19^. Like Sec14, Sfh3 also binds PI, and the crystal structure of the protein in a 1:1 complex with a PI molecule has been determined ^20^. Sfh3 also binds ergosterol (the major sterol in yeast), and ergosterol synthesis intermediates such as lanosterol and zymosterol ^21^. Moreover, *in vitro* assays showed that Sfh3 can transfer PI and sterols individually between membranes ^11, 21^. To date, the exact cellular function of Sfh3 and why it must recognize sterols and PI remain unknown.

In vegetatively growing yeast, Sfh3 is cytosolic but also localized to a subpopulation of LD^13–14, 22–24^. These structures, composed of a core of neutral lipids (triacylglycerols (TAGs), sterol esters) covered by a lipid monolayer, originate from the ER and serve as cellular energy stores^25^. Recent studies have shown that the Sfh3 localization to LDs depends on its physical interaction with Ldo45, attached to the LD monolayer^13–14, 22, 24^. Together with Ldo16, Ldo45 contributes to the formation of contact sites between LDs and vacuoles, facilitating microlipophagy—a process in which LDs are engulfed and digested within the vacuole to release fatty acyl chains when yeast enter the stationary phase ^14^ ^13, 22^ ^24^. However, Sfh3, despite its association with Ldo45, appears to be dispensable for this process.

Bankaitis and co-workers also identified a functional link between Sfh3 and LDs when examining the role of Sfh3 in diploid yeast cells exiting the mitotic cycle and sporulating in response to nutrient starvation^15^. During this process, four haploid nuclei are produced within the mother cell, and each nucleus is packaged within a prospore membrane, whose formation begins with the coalescence of post-Golgi vesicles and continues through an expansion step that remains poorly understood. It was found that cells lacking Sfh3 showed accelerated TAG degradation within LDs, while Sfh3 overexpression suppressed TAG breakdown. It has thus been proposed that defects in spore formation observed in Sfh3-overexpressing yeast arise from insufficient degradation of neutral lipids stored in LDs that are needed for prospore expansion.

Another line of research has shown that Sfh3 provides yeast cells, including pathogenic yeasts such as *Candida albicans*, with resistance to antifungal drugs that inhibit the ergosterol biosynthesis pathway^16–19^. This pathway consists of 20 metabolic steps ensured by enzymes localized at the ER and at LDs^26^; Sfh3 provides yeast with resistance to azoles that inhibit lanosterol 14-α-demethylase (Erg11), and increases cellular lanosterol levels at the expense of ergosterol, leading to cellular membrane destabilization. However, the molecular mechanism by which Sfh3 confers this resistance remains poorly understood. Data indicate that Sfh3 mitigates lanosterol accumulation at the expense of ergosterol production in miconazole-treated yeast, and this depends on its capacity to recognize PI ^16,18^. However, Sfh3 is not required for ergosterol biosynthesis in untreated cells. In addition, recent data suggest that the association of Sfh3 with Ldo45 at the LD surface and its potential capacity to mobilize TAG and ergosterol ester from LDs are not required for its protective role against azole treatment^27^.

Last, it has been found that abnormal ergosterol biosynthesis due to the deletion of genes coding for Erg6, Erg2, or Erg5, enzymes involved in late-stage ergosterol biosynthesis, or exposure to miconazole, provides yeast with resistance to Aureobasidin A in an Sfh3-dependent manner ^28^ ^29^. Because Aureobasidin A inhibits Aur1, the enzyme that catalyzes the first step in complex sphingolipid synthesis from ceramide, these results suggest that Sfh3 may function at the interface between sphingolipid and ergosterol metabolic pathways.

Our limited understanding of Sfh3’s functional role in both normal and azole-treated cells stems in part from our incomplete understanding of how Sfh3 recognizes and transfers its lipid ligands. Notably, unlike Sec14 and other lipid exchangers, such as Osh4 and Osh6 in yeast^30–31^ from the ORP family, its lipid-exchange capacity has not been examined. Here, using dehydroergosterol (DHE), an ergosterol analog, we showed in vitro that Sfh3’s ergosterol transfer activity is not activated by PI and is severely inhibited by lanosterol and zymosterol. Next, we showed that Sfh3 transfers PI as efficiently as Sec14 in the presence of sterols. Unexpectedly, we discovered that Sfh3 can also transport PI4P and act as a sterol/PI4P exchanger in vitro, but likely not powerfully in cells. By examining Sfh3 mutants and the thermal stability of Sfh3 in complex with different lipid ligands, we defined how Sfh3 recognizes sterols, PI, and PI4P and its relative affinity for these ligands. This allowed us to establish that PI activates Sfh3’s lanosterol transfer activity via an exchange process that occurs selectively even in the presence of ergosterol. Finally, we showed that Sfh3 must function as a lanosterol/PI exchanger to confer resistance to miconazole in yeast. Collectively, these data suggest that Sfh3 is a sterol-modulated PI/PI4P transporter that exclusively operates PI/lanosterol exchange in azole-treated yeast cells.

## Result

### PI has a dual effect on Sfh3-mediated ergosterol transfer

As a first step toward understanding Sfh3 function, we investigated how it transfers ergosterol between membranes *in vitro* using DHE, a fluorescent ergosterol analog that differs only by an additional double bond. We mixed donor liposomes (L_A_, 200 µM lipids) composed of DOPC (di-oleoyl-phosphocholine) and containing 5% DHE and 2.5% of a second fluorescent lipid, DNS-PE (1,2-dioleoyl-*sn*-glycero-3-phosphoethanolamine-N-(5-dimethylamino-1-naphthalenesulfonyl)), and acceptor liposomes composed only of DOPC (L_B_, 600 µM lipids). After injecting Sfh3 (1.5 or 4.5 µM), we measured a greater decrease in FRET between DHE and DNS-PE than after adding buffer alone, indicating that the protein actively transferred DHE from L_A_ to L_B_ liposomes. To confirm this, we used a positive control, cSTD3, which encompasses the sterol-binding START domain of STARD3. By normalizing the FRET signal, we established that the initial DHE transfer rate of Sfh3, measured at 1.5 µM or 4.5 µM (0.16–027 DHE *per* min *per* protein), was lower than that of cSTD3 (1.31) measured at 1.5 µM **(Fig. 1A, C)**. However, unlike cSTD3, we observed an acceleration in DHE transfer mediated by Sfh3 a few minutes after injection. This may reflect a transition of the Sfh3 pool, purified from *E. coli* in an apo, dimeric state, to a lipid-bound, monomeric state^20^, which could be more active. To test this, we mixed Sfh3 with L_A_ liposomes for one hour to promote its transition to a monomeric, DHE-loaded state and added this mix to L_B_ liposomes to initiate sterol transfer. We measured a transfer rate five times higher (1.5 DHE *per* min *per* protein) than when Sfh3 was not pre-mixed **(Fig. 1A, B)**, and observed no acceleration. Consequently, we pre-mixed Sfh3 with donor liposomes containing lipid ligands in all subsequent transfer assays to ensure the protein was in its most active state (unless otherwise stated).

**Figure 1.**
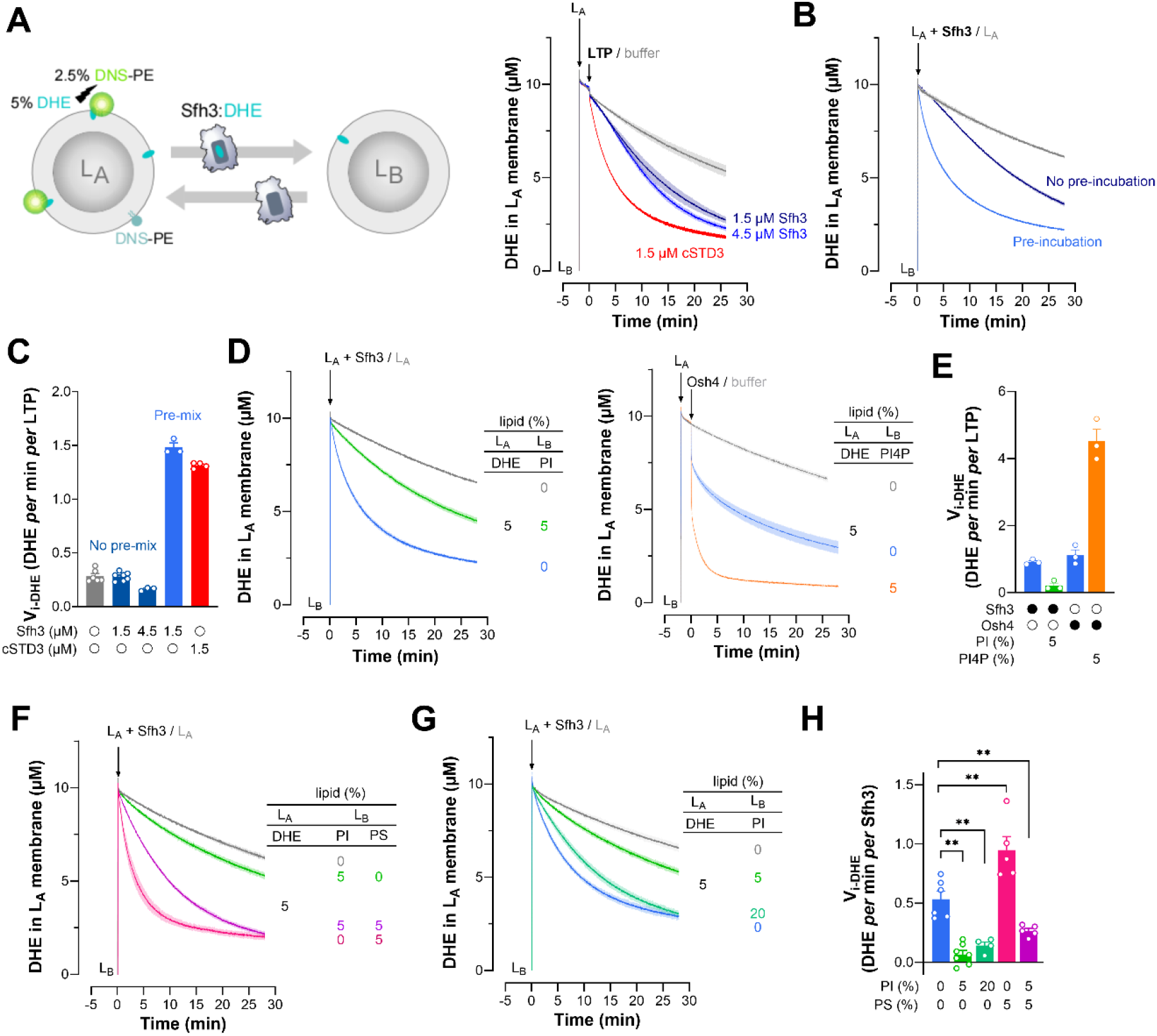
Sfh3-mediated DHE transfer is modulated by PI and by membrane negative charge. **(A)** DHE transfer assays. L_A_ liposomes composed of DOPC/DHE/DNS-PE (92.5:5:2.5, 200 µM total lipids) were mixed with L_B_ liposomes, made only of DOPC (600 µM), in HK buffer at 30 °C under constant stirring. Two minutes later, a small volume of Sfh3 or cSTD3 stock solution was added (1.5 or 4.5 µM, final concentration). Control experiments were performed by injecting a similar volume of HK buffer to measure spontaneous DHE transfer. Fluorescence was measured at λ_em_ = 525 nm upon excitation at λ_ex_ = 310 nm. The signal was normalized to the amount of DHE in the L_A_ membrane. Each trace is the mean ± s.e.m. of several kinetics recorded in independent experiments (n = 3–4). **(B)** Sfh3-mediated DHE transfer was measured as in (A) or by pre-mixing the protein with L_A_ liposomes for 1 h before adding the mixture to a cuvette containing L_B_ liposomes (red curve, pre-mixture). In each case, the final concentrations of protein and liposomes were the same (200 µM L_A_ liposomes, 600 µM L_B_ liposomes, 1.5 µM Sfh3). Control experiments were carried out without Sfh3. Each trace is the mean ± s.e.m. of several kinetics (with Sfh3, n = 3; without Sfh3, n = 2). **(C)** Initial DHE transfer velocities (V_i-DHE_) determined from (A) and (B) with Sfh3, pre-mixed or not with L_A_ liposomes, or cSTD3. Spontaneous DHE transfer was subtracted. Data are shown as the mean ± s.e.m with individual data points (with Sfh3, n = 3–7; with cSTD3, n = 4). **(D)** Comparative analysis of the influence of PI and PI4P on Sfh3- and Osh4-mediated DHE transfer, respectively. With Sfh3, experiments were performed as in (B), using L_B_ liposomes either composed solely of DOPC or containing 5% PI. Osh4 was added directly to L_A_ liposomes composed of DOPC/DHE/DNS-PE (92.5:5:2.5, 200 µM total lipids) and L_B_ liposomes (600 µM) composed of DOPC, with or without 5% brain PI4P. Control experiments were performed without LTP, using L_B_ liposomes purely composed of DOPC. Each trace corresponds to the mean ± s.e.m. of several kinetics (n = 3). **(E)** Initial DHE transfer velocities determined from (D). Spontaneous DHE transfer was subtracted. Data are shown as the means ± s.e.m. with individual data points (n = 3). **(F)** Comparative analysis of the influence of PS and PI on Sfh3-mediated DHE transfer. Experiments were performed as in (B) with L_B_ liposomes only made of DOPC or containing 5% PI, 5% PS, or both. Control experiments were performed without Sfh3 using L_B_ liposomes made only of DOPC. Each trace corresponds to the mean ± s.e.m. of several kinetics (with Sfh3, n = 3–5; without Sfh3, n = 2). **(G)** Impact of PI levels on Sfh3-mediated DHE transfer. Experiments were performed as in (B) with L_B_ liposomes only made of DOPC or containing 5 or 20% PI. Each trace corresponds to the mean ± s.e.m. of several kinetics (with Sfh3, n = 3–4; without Sfh3, n = 3) **(H)** Initial DHE transfer velocities determined from (F) and (G) as a function of the lipid composition of L_B_ membranes. Spontaneous DHE transfer was subtracted. Data are shown as the means ± s.e.m. with individual data points (n = 4–7). Unpaired Mann–Whitney U test (**p < 0.01).

Next, we investigated how Sfh3 exchanges DHE with PI between membranes. Based on our knowledge of lipid exchange processes mediated by ORPs, and notably the sterol/PI4P exchanger Osh4 ^31–32^, we hypothesized that Sfh3 would transfer DHE more rapidly to liposomes that contain PI than to pure DOPC liposomes. We found the opposite: the DHE transfer rate measured with Sfh3 was reduced 4-fold when L_B_ liposomes included 5% soy PI (hereafter called PI; otherwise specified) **(Fig. 1D, E)**. In comparison, we found that Osh4 transferred DHE 4-fold faster to L_B_ liposomes doped with 5% PI4P. Under conditions extensively used to study Osh4 activity ^31^, with L_A_ and L_B_ liposomes at a 1:1 ratio (200 µM lipid each), we confirmed that Sfh3-mediated DHE transfer is inhibited by PI and that Osh4-mediated DHE transfer was strongly activated by PI4P with an expected transfer rate of almost 20 DHE *per* min *per* protein^31^ **(Fig. S1A, B).** Thus, contrary to Osh4, Sfh3 is not an ergosterol transporter whose activity is boosted by its second lipid ligand *via* an exchange process.

We further investigated whether PI inhibition of Sfh3 might result from its retention on the L_B_ membrane through nonspecific electrostatic interactions with PI, thereby impairing its ability to rapidly cycle between liposomes. We therefore measured how Sfh3 transferred DHE to L_B_ liposomes containing either 5% PI or another anionic lipid, phosphatidylserine (PS). We found that, whereas Sfh3’s transfer activity was reduced in the presence of PI, it was higher in the presence of PS. Interestingly, we measured an intermediate rate with L_B_ liposomes containing 5% of both lipids, suggesting that PS counteracts PI inhibitory effect on Sfh3’s sterol transfer activity **(Fig. 1F, H)**. Flotation assays revealed that Sfh3 binds weakly to DOPC membranes, whether or not they are doped with 5% DHE, and binds slightly more, but similarly, to membranes containing either 5% PI or PS **(Fig. S1C)**. We concluded that small amounts of anionic lipids may accelerate Sfh3-mediated DHE delivery to membranes, but that PI specifically inhibits this process, likely because it is specifically recognized by Sfh3.

Since yeast membranes contain a high level of PI (20–25% of total lipids)^33–34^, we tested how Sfh3 transferred DHE in the presence of L_B_ liposomes containing 20% PI. We found that it was more active than with L_B_ liposomes containing 5% PI, but still less than with PI-free L_B_ liposomes **(Fig. 1G, H)**. Soy PI mostly consists of 16:0/18:2-PI (70%) and thus cannot completely mimic the diversity of PI species found in yeast, whose prominent ones are 16:0/16:1-PI and 16:0/18:1-PI^33^. We therefore repeated our measurements with L_B_ liposomes containing 0%, 5%, or 20% yeast PI and confirmed that, despite slight quantitative differences, Sfh3 DHE-transfer activity was reduced in the presence of PI **(Fig. S1D)**. We also examined Sfh3’s ability to transfer DHE to L_B_ liposomes containing 5% PI and increasing amounts of PS and found that it was severely impaired when the PS fraction reached 20% **(Fig. S1E, F)**. Flotation assays showed that Sfh3 bound strongly to membranes with 20% PS and more weakly to membranes with the same amount of PI **(Fig. S1G)**. Overall, these data suggest that the ergosterol transfer activity of Sfh3 is not activated by PI in yeast cells and can be rather inhibited to an extent that depends on the density of PI, which modulates the surface negative charge of the membrane, as well as its acyl-chain composition, which likely affects how PI is recognized by Sfh3.

### Lanosterol and zymosterol block ergosterol transfer by Sfh3

Lanosterol, an intermediate of the ergosterol biosynthesis pathway **(Fig. 2A)**, was found to be an Sfh3 ligand^21^. To compare lanosterol with ergosterol, we measured how Sfh3 exchanged DHE between L_A_ doped with 5% DHE and L_B_ liposomes containing or not containing 5% ergosterol or lanosterol. Strikingly, its transfer activity, unchanged in the presence of ergosterol, was abrogated by lanosterol **(Fig. 2B, C)**. The same inhibitory effect was seen with tenfold less lanosterol (0.5 %). Moreover, another ergosterol intermediate, zymosterol, could block DHE transfer. In comparison, the ability of cSTD3 to transfer DHE remained unaffected, regardless of the sterol species incorporated into the L_B_ membranes **(Fig. 2B, C)**. This suggests that Sfh3’s binding pocket can specifically host lanosterol or zymosterol, possibly with an affinity higher than for ergosterol.

**Figure 2.**
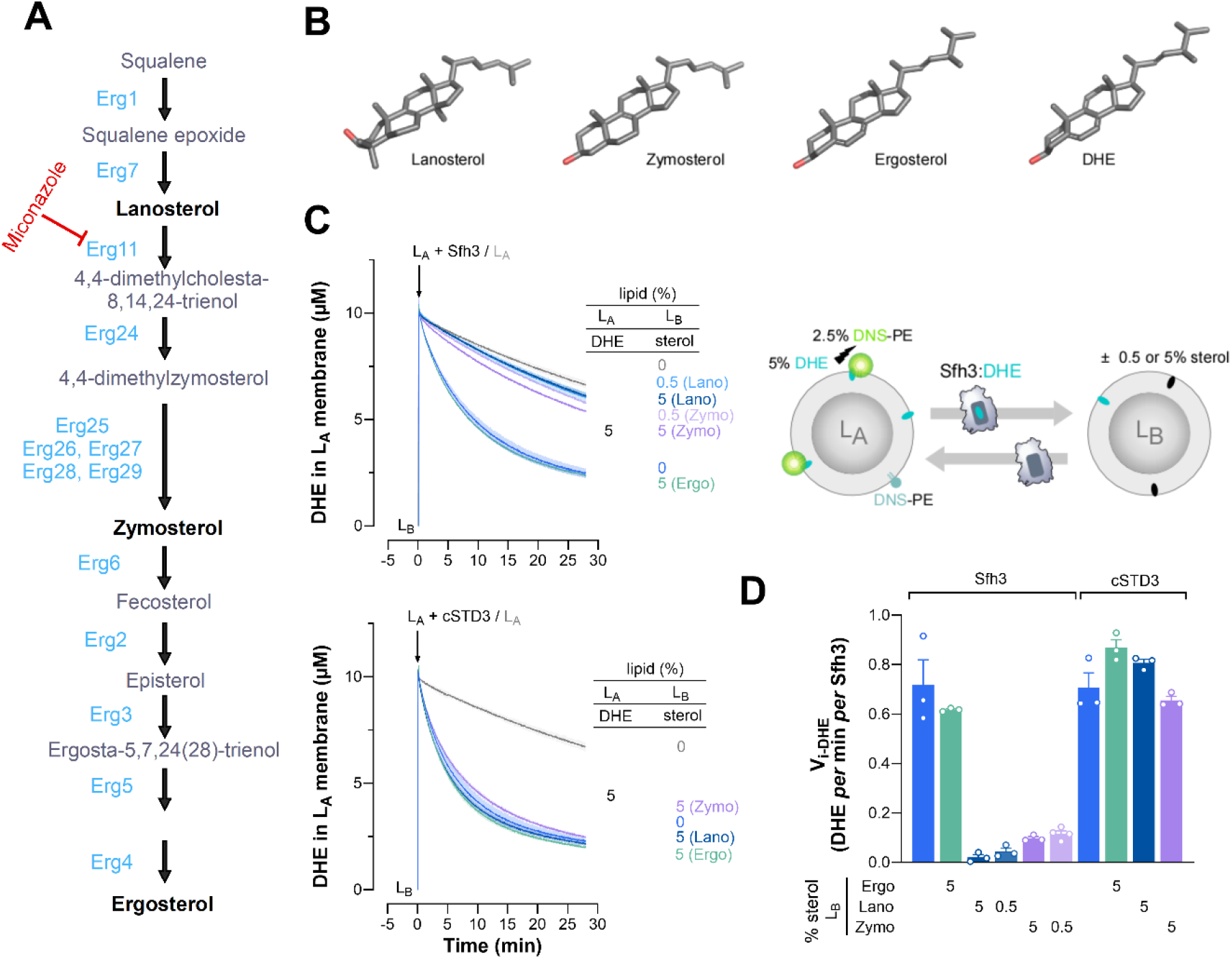
Lanosterol and zymosterol inhibit the DHE transfer activity of Sfh3. **(A)** Ergosterol biosynthesis pathway in *S. cerevisiae*. Ergosterol intermediates are shown in grey, and enzyme names are in blue. Miconazole inhibits ergosterol synthesis by targeting lanosterol 14α-demethylase, encoded by the ERG11 gene. This inhibition provokes an increase in lanosterol levels at the expense of ergosterol in yeast cells. **(B)** Three-dimensional structure of lanosterol, zymosterol, ergosterol, and DHE. Carbon atoms are in grey, oxygen atoms are in red, and hydrogen atoms are not represented. **(C)** DHE transfer assay. Sfh3 or cSTD3 was incubated with L_A_ liposomes composed of DOPC/DHE/DNS-PE (92.5:5:2.5) for 1 h at 30 °C, and the mixture was added to L_B_ liposomes, made only of DOPC or additionally containing 0.5 or 5% ergosterol (Ergo), lanosterol (Lano), or zymosterol (Zymo). The final concentrations of LTP, L_A,_ and L_B_ liposomes were 1.5, 200, and 600 µM, respectively. Control experiments were done without LTP. Each trace is the mean ± s.e.m. of 3–4 kinetics. **(D)** Initial DHE transfer velocities determined from (C) with Sfh3 and cSTD3 as a function of the sterol content of L_B_ liposomes. Spontaneous DHE transfer was subtracted. Data are represented as mean ± s.e.m. (n = 3–4) with single data points.

### Sfh3 transfers PI between membranes in a sterol-dependent manner

Next, we measured in real time the PI transfer activity of Sfh3. We incubated Sfh3 with L_A_ liposomes enriched with 20% PI and 2% Rhod-PE, then added them to L_B_ liposomes coated with a PI-4 kinase (L_B_-PI4K). PI transfer from L_A_ to L_B_ membranes was quantified by measuring the phosphorylation of PI to PI4P by the kinase in the presence of ATP, using the sensor NBD-PH_FAPP_^35^ whose fluorescence increases only when PI4P appears in L_B_ membranes. Sfh3 promoted PI4P accumulation, albeit to a lesser extent than Sec14, which was used as a positive control, suggesting that Sfh3 is a less active PI-transfer protein than Sec14 **(Fig. 3A)**. We reasoned that, compared with Sec14, which could exchange PI for PC in our assay, the Sfh3 PI transfer activity was measured in the absence of a counterligand (i.e., sterol), which could explain its lower activity. Therefore, we repeated the experiments using L_B_-PI4K liposomes containing 10% DHE and measured that Sfh3 promoted a higher PI4P production, comparable to that observed with Sec14. We observed a weaker effect when 10% DHE was incorporated into L_A_ membranes, and obtained an intermediate result when sterol was included in both L_A_ and L_B_-PI4K liposomes. In parallel, we analyzed how Sec14 promoted PI4P synthesis using the same liposome combinations and measured nearly identical activity **(Fig. 3B, C)**, suggesting that this LTP is not impacted by sterol, as it does not recognize it.

**Figure 3.**
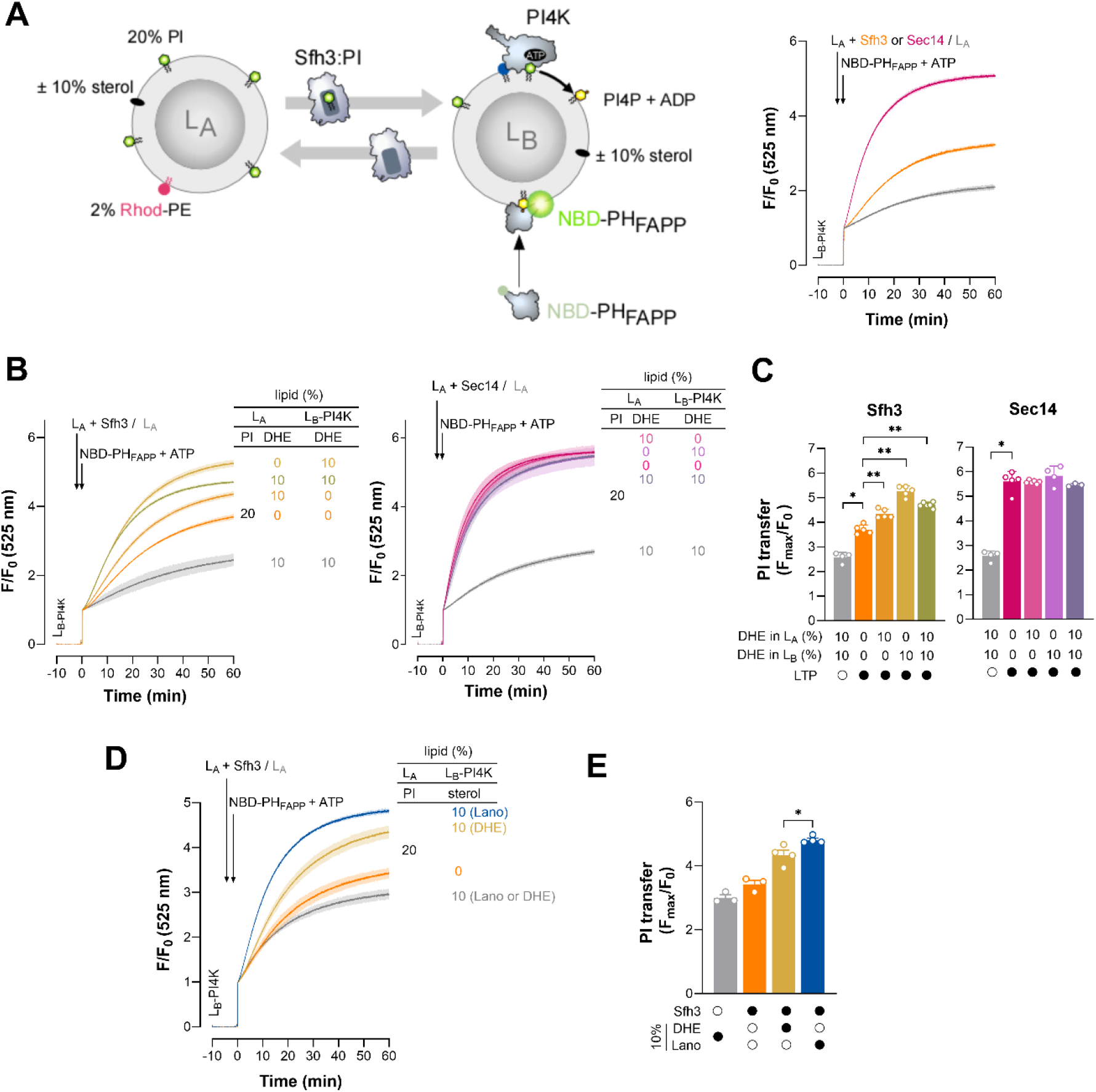
Sfh3-mediated PI transfer is modulated by the sterol levels in the membranes. **(A)** PI-transfer assay. L_A_ liposomes composed of DOPC/PI/Rhod-PE (78:20:2) were incubated with or without Sfh3 or Sec14 for 1 h at 30 °C and then added to L_B_-PI4K liposomes, composed of DOPC/MPB-PE (97:3) and coated with PI-4 kinase (P/L = 1/400). Final concentration: Sfh3 or Sec14, 200 nM; L_A_ and L_B_ liposomes, 200 µM lipids each. Thirty seconds later, NBD-PH_FAPP_ (250 nM) and ATP (100 µM) were injected together. Fluorescence was measured at 525 nm (λ_ex_ = 460 nm) at 30 °C under constant stirring and normalized to the signal (F_0_) measured immediately after the last injection. Each trace is the mean ± s.e.m. of several kinetic traces (with Sfh3 or Sec14, n = 3; no LTP, n = 2). **(B)** The same experiments as in (A) were performed with Sfh3 and Sec14 with different combinations of L_A_ and L_B_-PI4K liposomes, containing or not 10% DHE. Control experiments were carried out without LTP, with liposomes containing 10% DHE. Mean ± s.e.m (with Sfh3, n = 5–6; with Sec14, n = 3–6, without LTP, n = 2). **(C)** PI transfer rate corresponding to the F_max_ signal measured at the end of the kinetics shown in (B), normalized to F_0_, carried out with Sfh3 and Sec14, as a function of the DHE content of liposomes. Data are represented as mean ± s.e.m. (n = 4–6) with single data points. Unpaired Mann–Whitney U test (* *p* < 0.05, \*\**p* < 0.01). **(D)** The same experiments as in (A) were performed with Sfh3 with L_B_-PI4K liposomes, containing or not 10% DHE or 10% lanosterol. Control experiments were carried out without LTP using L_B_-PI4K liposomes containing either 10% DHE or lanosterol. Data are represented as mean ± s.e.m (with Sfh3, n = 3–4, without Sfh3, n = 3). **(E)** PI transfer rates extracted from kinetics shown in (D) as a function of the sterol content of L_B_-PI4K liposomes. Data are represented as mean ± s.e.m. (n = 3–4) with individual data points. Unpaired Mann–Whitney U test (\*\**p* < 0.01).

Next, we examined how Sfh3 transferred PI to L_B_-PI4K liposomes either lacking sterol or containing 10% DHE or lanosterol. We found that Sfh3 promoted significant PI4P accumulation in the presence of lanosterol, slightly higher than in the presence of DHE (**Fig. 3D, E)**. A similar result was obtained with zymosterol (**Fig. S2A**). Thus, sterol in the L_B_-PI4K membrane enhances Sfh3-mediated PI4P synthesis, likely because Sfh3 exchanges sterol for PI, thereby facilitating net delivery of PI into the L_B_ liposomes. Eventually, we found that Sfh3 transferred yeast PI between L_A_ and L_B_-PI4K liposomes enriched with 10% DHE **(Fig. S2B)**. These data suggest that Sfh3 can properly transfer PI between sterol-rich membranes in yeast.

### Sfh3 has a PI4P transfer activity

Next, we explored whether the lower fluorescence signals measured in the kinase assay with Sfh3, compared to Sec14, might also result from Sfh3 transferring PI4P from L_B_-PI4K to L_A_ liposomes once this lipid is synthesized. Indeed, such a transfer would cause the translocation of NBD-PH_FAPP_ to L_A_ liposomes and the quenching of its fluorescence by Rhod-PE. To do so, we tested whether Sfh3 and Sec14 moved PI4P from L_B_-PI4K to L_A_ liposomes under conditions in which 25% of the accessible PI initially in L_A_ liposomes would have been transferred into L_B_-PI4K liposomes and converted into PI4P. Each LTP was added to L_B_ liposomes composed of PC/PI/PI4P/Rhod-PE (85.5:7.5:5:2, mol/mol) mixed with L_A_ liposomes composed of PC/PI (92.5:7.5) and NBD-PH_FAPP_ (see Materials and Methods for details). Adding Sec14 did not provoke dequenching of the NBD signal, indicating it did not transport PI4P. In contrast, adding Sfh3 triggered dequenching, revealing that it can transport PI4P. A slightly lower transfer was observed if L_B_ liposomes contained 10% DHE or zymosterol, whereas a strong reduction was seen with L_B_ liposomes enriched with 10% lanosterol **(Fig. 4A)**. This suggested that lanosterol blocked the transfer of PI4P to L_A_ liposomes and could explain why a higher PI4P accumulation was seen when L_B_-PI4K contained 10% lanosterol instead of DHE **(Fig. 3D, E)**. These data suggest that Sfh3 promotes higher PI4P enrichment in sterol-rich L_B_-PI4K membranes by delivering PI more efficiently and extracting less PI4P.

**Figure 4.**
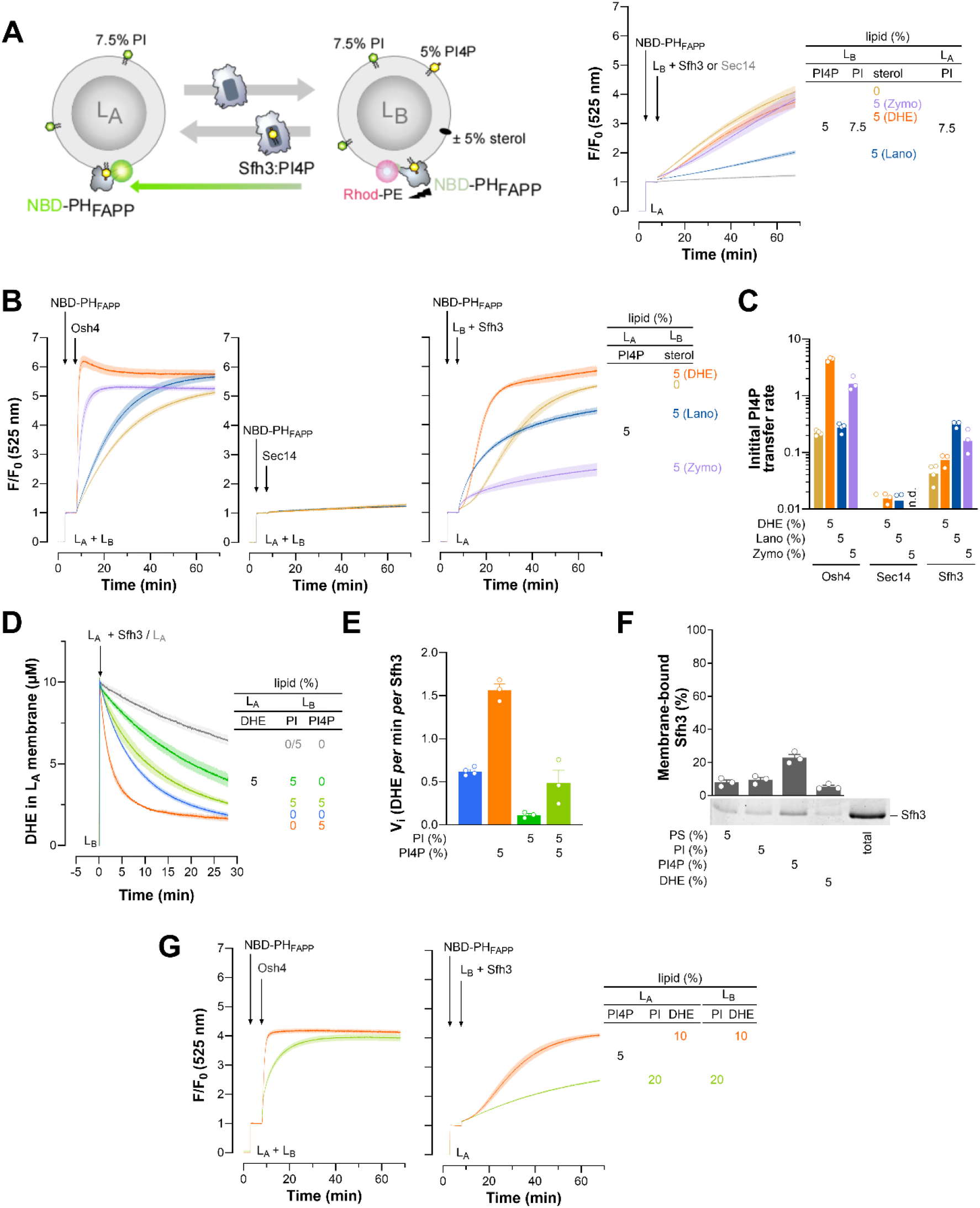
Sfh3 exhibits PI4P transfer activity. **(A)** PI4P-transfer assay. L_B_ liposomes containing 7.5% PI, 5% 18:1/18:1-PI4P, 2% Rhod-PE, doped or not with sterol (DHE, lanosterol, or zymosterol), were incubated with Sfh3 or Sec14 for 1 h at 30 °C and then added to L_A_ liposomes containing 7.5% PI in the presence of NBD-PH_FAPP_. Final LTP concentration: 200 nM; L_A_ and L_B_ liposomes: 200 µM lipids each; NBD-PH_FAPP_: 250 nM. Fluorescence was measured at 525 nm (λ_ex_ = 460 nm) at 30 °C upon constant stirring and normalized to the signal (F_0_) measured immediately after the last injection. Each trace is the mean ± s.e.m. of several kinetics (n = 3–5). **(B)** Sfh3 was incubated with L_B_ liposomes made of DOPC or additionally containing 5% DHE, lanosterol, or zymosterol for 1 h; the mixture was then added to L_A_ liposomes containing 5% PI4P, pre-mixed with NBD-PH_FAPP_. Osh4 or Sec14 was added directly to L_A_ and L_B_ liposomes in the presence of NBD-PH_FAPP._ Final LTP concentration, 200 nM; L_A_ and L_B_ liposomes, 200 µM lipids each; NBD-PH_FAPP_, 250 nM. Each trace is the mean ± s.e.m. of 3–4 kinetics. **(C)** Initial PI4P transfer rate determined from (B) with Sfh3 or Osh4, as a function of the lipid composition of L_B_ liposomes. Data are represented as mean ± s.e.m. (n = 2–4) with single data points **(D)** DHE transfer assay. Sfh3 was incubated with L_A_ liposomes composed of DOPC/DHE/DNS-PE (92.5:5:2.5, 200 µM total lipids) for 1 h, and the mixture was injected into a cuvette containing L_B_ liposomes made of DOPC, containing or not 5% PI and/or 18:1/18:1-PI4P. Final concentrations: L_A_ liposomes, 200 µM; L_B_ liposomes, 600 µM; Sfh3, 1.5 µM. A control experiment was performed without Sfh3. Each trace is the mean ± s.e.m. of several kinetics (n = 3–4). **(E)** Initial DHE-transfer velocities determined from (D) as a function of the lipid composition of L_B_ membranes. Spontaneous DHE transfer was subtracted. Data are represented as mean ± s.e.m. (n = 3–4) with single data points. **(F)** Flotation assay. Sfh3 was incubated with liposomes made of DOPC and containing 5% PI, PS, PI4P, or DHE. Data are represented as mean ± s.e.m. (n = 3) with single data points. **(G)** Sfh3 was pre-incubated with L_B_ liposomes containing 10% DHE, and the mixture was added to L_A_ liposomes containing 10% DHE and 2% Rhod-PE in the presence of NBD-PH_FAPP_. Osh4 was added directly to L_A_ and L_B_ liposomes pre-mixed with NBD-PH_FAPP._ Measurements were repeated with L_A_ and L_B_ liposomes incorporating PI (20%) instead of DHE. Final concentration: Sfh3 or Osh4, 200 nM; L_A_ and L_B_ liposomes, 200 µM lipids each; NBD-PH_FAPP_, 250 nM. Each trace is the mean ± s.e.m. of several kinetics (n = 3–4).

Because Sfh3 can transfer both DHE and PI4P, we investigated whether it could act as a sterol/PI4P exchanger, similar to Osh4. To test this, we pre-incubated Sfh3 with L_B_ liposomes composed of DOPC and then added it to L_A_ liposomes enriched with 5% PI4P and 2% Rhod-PE, in the presence of NBD-PH_FAPP_. In parallel experiments, we measured the PI4P transfer activity of Osh4 or Sec14 (as a negative control) by directly adding these LTPs to the two liposome populations. Except for Sec14, we observed NBD dequenching, which allowed us to calculate an initial PI4P transfer rate for Sfh3 that was 5.5-fold lower than that of Osh4 **(Fig. 4B, C)**. The dequenching kinetics for Sfh3 were sigmoidal, suggesting that Sfh3, pre-incubated with ligand-free L_B_ liposomes, shifted from a dimeric to a monomeric state upon binding and transporting PI4P originating from L_A_ liposomes. When L_B_ liposomes were doped with 5% DHE, Sfh3 transferred PI4P 1.7-fold faster, possibly through DHE/PI4P exchange and/or because Sfh3 was pre-bound to DHE in a monomeric state. In comparison, Osh4 activity increased 20-fold and was 60-fold higher than that of Sfh3 **(Fig. 4B, C)**. We also repeated the assay with L_B_ liposomes enriched with 5% lanosterol. Osh4-mediated PI4P transfer was barely accelerated, suggesting that Osh4 does not recognize lanosterol or exchange it for PI4P. In contrast, the initial rate of Sfh3-mediated PI4P transfer increased nearly 8-fold compared to the rate observed with sterol-free L_B_ liposomes. However, the transport process rapidly slowed, suggesting that lanosterol accumulation in L_A_ liposomes inhibits Sfh3-mediated PI4P removal **(Fig. 4B)**. Eventually, we observed that zymosterol accelerated Osh4-mediated PI4P transfer activity but abolished Sfh3 activity. Together, these results suggest that Sfh3 can transfer PI4P and potentially exchange it for sterols, albeit much less efficiently than Osh4.

To further assess whether Sfh3 could function as a sterol/PI4P exchanger, we measured its ability to transfer DHE from L_A_ liposomes to L_B_ liposomes containing or not 5% PI4P. We observed increased DHE transfer in the presence of PI4P **(Fig. 4D, E)**. In comparison, when L_B_ liposomes contained 5% PI, Sfh3 DHE transfer activity was inhibited, as previously observed. We measured intermediate activity when L_B_ liposomes contained both 5% PI and PI4P, suggesting that PI competes with PI4P and sterols for binding to Sfh3’s lipid-binding pocket. Using flotation assays, we found that Sfh3 binds more strongly to membranes containing 5% PI4P than to those containing 5% PI **(Fig. 4F)**. The fact that Sfh3 transfers DHE more efficiently to L_B_ liposomes containing PI4P instead of PI, despite PI4P retaining more Sfh3 on these liposomes than PI, supports the idea that Sfh3 efficiently exchanges DHE for PI4P.

Finally, we evaluated how rapidly Sfh3 and Osh4 transfer PI4P between membranes rich in either sterols or PI, mimicking yeast endomembranes. As previously observed^31^, Osh4 efficiently transferred PI4P from L_A_ to L_B_ liposomes enriched with 10% DHE, enabling rapid PI4P equilibration between membranes within seconds **(Fig. 4G)**. In contrast, Sfh3 was much less active. When we repeated the experiment with liposomes enriched with 20% PI, Osh4 retained substantial activity, whereas Sfh3’s activity was almost abolished. These data suggest that Sfh3 is unlikely to primarily function as a robust ergosterol/PI4P exchanger like Osh4 in yeast cells.

### Analysis of the DHE and PI transfer activities of Sfh3 mutants

To further analyze Sfh3 function, we carried out a structure-function analysis by examining how various Sfh3 mutants transfer sterol and PI in our assays (see Sfh3 sequence in **Fig. S3A**). Some of these mutants have been tested *in vitro*, but in assays that analyzed sterol and PI transfer separately, with no or low temporal resolution^11, 15, 21^. To select the mutants, we also considered the structure of Sfh3 bound to PI in an open conformation (PDB ID 4J7Q^20^) and in a closed conformation, obtained by homology modeling with the PI-bound closed form of Sfh1 and molecular docking ^20^. We purified these mutants and confirmed that they were well folded using circular dichroism **(Fig. S3B–D)**. We then assessed their DHE transfer activity in the presence or absence of 5% PI (as in **Fig. 1D**), and their PI transfer activity in the presence of lanosterol (as in **Fig. 3D**; since lanosterol blocks PI4P transfer, this allowed us to unambiguously test how mutations in Sfh3 affect its PI transfer activity).

We first tested Sfh3 mutants in which the E235 and R107 residues, whose side chains form hydrogen bonds with the 2-OH and 4-OH groups of the inositol ring of PI in the open structure of Sfh3, respectively ^20^, were mutated to alanine (**Fig.5A**). The DHE transfer activity of Sfh3^E235A^ and Sfh3^R107A^ was similar to that of Sfh3 WT but was less sensitive to PI (**Fig.5B** and **Fig.S4**). Both mutants transferred PI almost as efficiently as Sfh3 WT (**Fig. 5C** and **Fig. S5A**). We then tested a double mutant, Sfh3^E235A/K267A^, in which we also mutated K267, which forms a hydrogen bond with the phosphate group at position 1 of inositol^20^. This mutant showed markedly elevated DHE transfer activity, which PI could further accelerate, but no PI transfer activity. This result aligns with previous findings showing no PI transfer activity for this mutant, but contradicts those suggesting that it also lacks sterol transfer capacity^11^.

**Figure 5.**
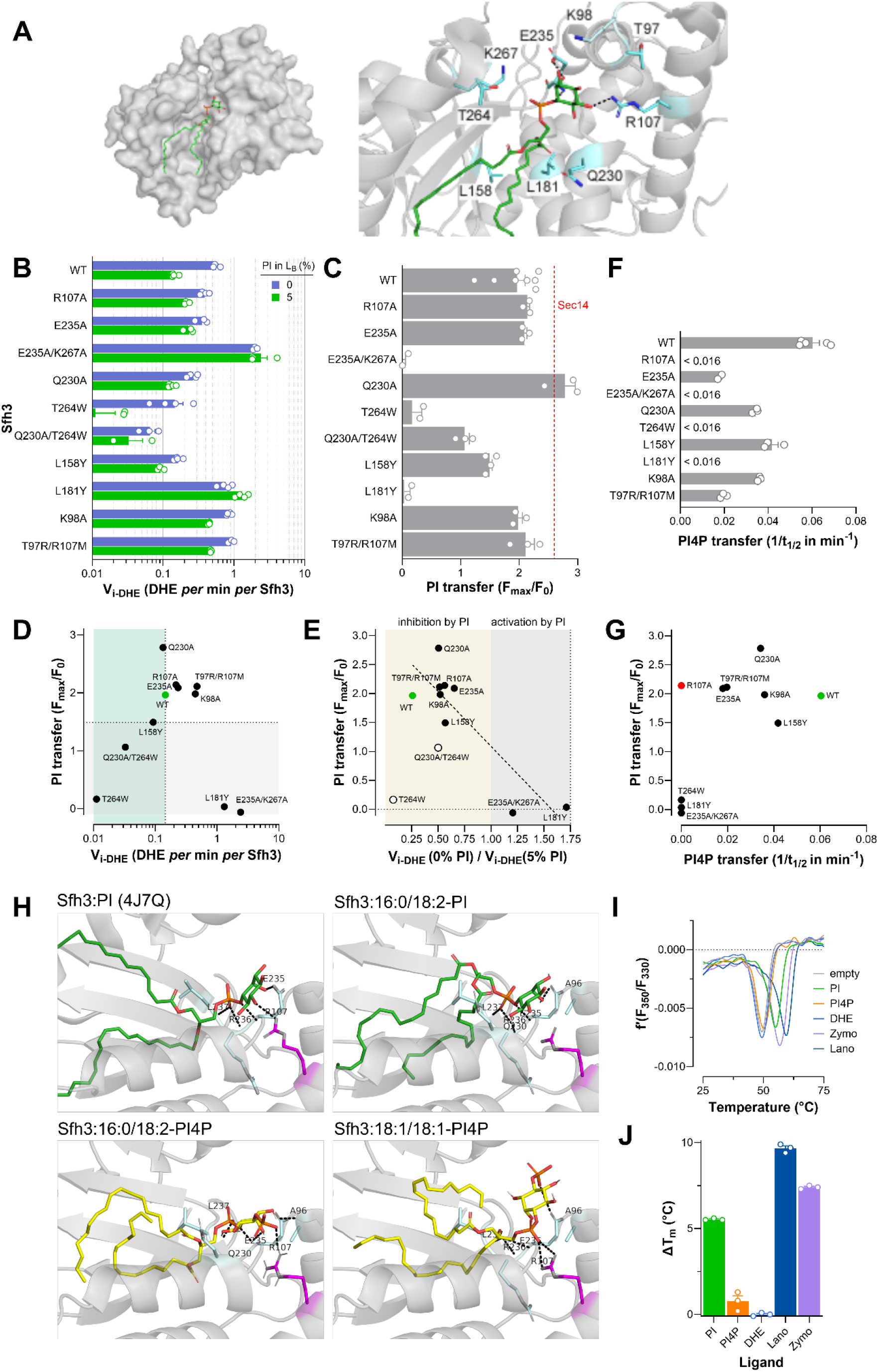
Analysis of the DHE, PI, and PI4P transfer activities of Sfh3 mutants. **(A)** Crystal structure of the Sfh3:PI complex (PDB 4J7Q, chain A) with the protein represented as a surface and the PI molecule drawn as sticks. The residues forming hydrogen bonds (dashed black lines) with PI are shown and labeled in a close-up view of the structure. **(B)** Initial DHE transfer velocities were measured between L_A_ liposomes composed of DOPC/DHE/DNS-PE (92.5:5:2.5), pre-incubated with Sfh3 WT or mutants, and L_B_ liposomes only made of DOPC (blue bars) or additionally containing 5% PI (green bars, see also **Fig.S4**). Final concentration: Sfh3 WT or mutants, 1.5 µM; L_A_ and L_B_ liposomes, 200 and 600 µM lipids, respectively. Spontaneous DHE transfer velocities were measured without Sfh3 in both conditions and subtracted from velocities determined with proteins. Data are represented as mean ± s.e.m. (n = 3–4) with single data points. **(C)** PI transfer activity of Sec14, Sfh3 WT or mutants (200 nM) measured with L_A_ liposomes (200 µM lipids) composed of DOPC/PI/Rhod-PE (78:20:2) and L_B_-PI4K liposomes (200 µM lipid) composed of DOPC/lanosterol/MPB-PE (87:10:3), coated with PI4K at P/L =1/400. Control experiments were performed without LTP. Data are shown as mean ± s.e.m. (n = 3–9) with individual data points. The mean PI transfer activity of Sec14 is indicated with a red dashed line (n = 3). See also **Fig. S5A**. **(D)** PI transfer rates of Sfh3 WT and mutants are shown as a function of their initial DHE transfer velocities measured in the presence of 5% PI. **(E)** PI transfer rates of Sfh3 WT and mutants are shown as a function of the ratio of initial DHE transfer velocities measured for each construct in the presence and absence of 5% PI. Data points, except those corresponding to Sfh3^T264W^ and Sfh3^Q230A/T264W^ (open circles), were fitted with a linear equation (dashed line). **(F)** PI4P transfer rate of Sfh3 WT or Sfh3 mutant (200 nM), expressed in 1/t_1/2,_ measured between L_A_ liposomes (200 µM lipids) containing 5% PI4P and an equivalent amount of L_B_ liposomes made only of DOPC. Data are represented as mean ± s.e.m. (n = 2–3) with single data points. See also **Fig.S5B** **(G)** PI transfer rates of Sfh3 WT and mutants are plotted against their PI4P transfer rates. **(H)** Crystal structure of the Sfh3-PI complex and docked models of Sfh3 with 16:0/18:2-PI, 16:0/18:2-PI4, and 18:1/18:1-PI4P. The lipid is drawn as sticks, and the side chains forming hydrogen bonds are shown and labeled; hydrogen bonds are drawn as dashed black lines, with distances given in Table S1. Arg107 is colored magenta throughout. It forms hydrogen bonds with the inositol ring of PI in the Sfh3:PI crystal structure and with the head group of PI4P in both Sfh3:PI4P complexes. **(I)** Thermal stability of Sfh3 in an empty state or in complex with ligands. In a typical measurement, a sample containing 10 μM of protein in HK buffer was heated, and fluorescence (F) was measured at λ_em_ = 330 and 350 nm; the first derivative (f’) of the F_350_/F_330_ ratio gives access to the inflection point approximating the melting temperature (T_m_), reflecting the stability of the protein. **(J)** Difference in melting temperatures (ΔT_m_), derived from inflection points, between Sfh3 loaded with lipid ligands and empty Sfh3. Data are represented as mean ± s.e.m. (n = 3).

We then tested Sfh3^Q230A^, Sfh3^T264W^, and Sfh3^Q230A/T264W^, previously characterized by Bankaitis team^7^. We found that the Q230A mutant has a DHE transfer activity similar to that of Sfh3 WT, which can be inhibited by PI (**Fig. 5B** and **Fig. S4**). Remarkably, this mutant has a higher PI transfer capacity than Sfh3 WT, matching that of Sec14 used as a reference (**Fig. 5C** and **Fig. S5A**). Conversely, we found that substituting T264 with tryptophan abolished Sfh3’s PI and sterol transfer activities. Replacing the T264 residue with a bulky residue likely creates a steric clash that hinders accommodation of both lipids. Interestingly, analysis of the Sfh3^Q230A/T264W^ mutant indicates that the Q230A mutation partially reverses the deleterious effect of the T264W mutation on the PI transfer activity of Sfh3, possibly by enabling the protein to adopt an alternative closed conformation to host PI. We noted that our results did not align with previous ones, showing that Sfh3^Q230A^ is less efficient in transporting PI and sterol than Sfh3 WT, and contradicting the claims that the T264W mutation only abrogates the PI transfer activity of Sfh3^11, 15^.

Next, we tested the L158Y and L181Y mutations, designed to prevent Sfh3 from accommodating sterol in its pocket by replacing leucine with a bulky tyrosine residue^11^. The Sfh3^L158Y^ mutant has almost no sterol transfer activity, while its PI transfer activity is 25% less than that of Sfh3 WT (**Fig. 5B** and **C, Fig. S4** and **S5A**). In contrast, compared to Sfh3 WT, the Sfh3^L181Y^ mutant showed higher sterol transfer activity, which can be activated by PI, but no PI transfer activity. Overall, these data align with previous findings, showing that the L158Y and L181Y mutations respectively abrogate the sterol and PI transfer activities of Sfh3 in a rather specific manner^7^. Because L158 and L181 residues are adjacent, this suggests that introducing a tyrosine in the pocket may hinder the accommodation of the DHE or PI acyl chains, depending on its position.

Finally, we found that mutating the K98 residue, whose corresponding residue in Sfh1, Sfh2, and Sec14 is involved in PI recognition ^36–37^, to alanine weakly affects Sfh3 function. We also noted that the R107 residue in Sfh3, which contacts PI, has no equivalent in Sec14 and other Sfh proteins, where it is replaced by a non-anionic residue (methionine, asparagine, or glycine, **Fig. S3A**). In contrast, Sec14, Sfh1, and Sfh2 possess an arginine residue involved in PI recognition^36–37^ that is replaced by a threonine in Sfh3 (T97). We therefore tested an Sfh3^T97R/R107M^ mutant in which the T97 and R107 residues were replaced by their corresponding residues in Sec14 **(Fig. S3A)**, but found that it behaved like Sfh3 WT (**Fig. 5B and C, Fig. S4A and S5A**).

To provide a global analysis of our data, we plotted the PI transfer activity of Sfh3 WT and mutants against the DHE transfer activity of each construct in the presence of PI **(Fig.5D)**. This highlights five remarkable mutants: Sfh3^L181Y^ and Sfh3^E235A/K267A^ with the highest sterol transfer activity but no PI transfer activity, Sfh3^L158Y^ defective in sterol transfer but retaining a robust PI transfer activity, Sfh3^T264W^ defective in both sterol and PI transfer, and Sfh3^Q230A^ with a PI transfer activity superior to that of Sfh3 WT. We also plotted the PI transfer activity of Sfh3 WT and mutants against the ratio of DHE transfer rates measured for each construct in the absence and presence of PI (< 1, inhibition by PI, > 1 activation by PI, **Fig.5E**). We found a negative correlation (R^2^ = 0.76) between this ratio and the PI transfer capacity of Sfh3 WT and mutant (once excluded Sfh3^T26W^ and Sfh3^T264W/Q230A^ mutants from this analysis, as they are unable to transfer both PI and DHE). This confirms that Sfh3 inhibition of sterol transfer depends on the selective binding of PI by this LTP. It also suggests that when PI is not correctly hosted in Sfh3’s pocket, Sfh3 sterol transfer activity is activated because PI also provides a negative surface charge to the membrane, confirming data from **Fig. 1**.

### Analysis of the PI4P transfer activities of Sfh3 mutants

We further measured the capacity of a subset of Sfh3 mutants to transfer PI4P. We found that all these mutants were less efficient at transferring PI4P than Sfh3 WT, with Sfh3^R107A^, Sfh3^T264W^, Sfh3^L181Y^ and Sfh3^E235A/K267A^ being the most defective (1/_t1/2_ < 0.016 min^-1^, **Fig. 5F, Fig. S5B**). We plotted PI against PI4P transfer activities of these proteins **(Fig. 5G)**, which highlighted that the Sfh3^R107A^ mutant transfers PI as efficiently as Sfh3 WT but completely fails to transfer PI4P. However, an equivalent R107M mutation can be compensated by replacing T97 with arginine. Because the R107 residue is not conserved in Sec14 **(Fig. S3A)**, this suggests it is critical for PI4P recognition by Sfh3.

To better understand these results, we modeled how Sfh3 could recognize a PI4P molecule (with either 16:0/18:2 or 18:1/18:1 acyl chains) via molecular docking. The first PI4P species corresponds to what is primarily generated by PI4K in the PI transfer assay, whereas the second is used in the PI4P transfer assay. As a control, we also modeled how Sfh3 accommodates 16:0/18:2-PI. We first observed that, in the models with the 16:0/18:2 species, the lipid adopts a configuration similar to that of the PI molecule co-crystallized with Sfh3^20^ and is recognized by Sfh3 via a similar polar interaction network. Notably, E235 and L237 form hydrogen bonds with the 2-OH group of the inositol ring and an oxygen of the 1-phosphate group in both PI and PI4P. In the model with 16:0/18:2-PI, the R236 residue also forms a hydrogen bond with the 1-phosphate group of the lipid **(Fig. 5H, Table S2)**. We further observed that R107, which forms a hydrogen bond with the 4-OH group of the inositol ring of PI in the crystal structure, adopts the same orientation in the Sfh3:16:0/18:2-PI model but lies 3.42 Å from the head group, just beyond our hydrogen-bond criterion. Importantly, we found in our model of the Sfh3:PI4P complex that the R107 residue directly contacts an oxygen of the 4-phosphate group that distinguishes this lipid from PI (3.07 Å, **Fig. 1H, Table S2**). However, in our model of Sfh3 in complex with 18:1/18:1-PI4P, this lipid occupies a different position in the binding pocket, with E235, R236, and L237 forming hydrogen bonds with the carbonyl group of the sn-2 acyl chain. Nevertheless, the R107 residue remains engaged with the 1-phosphate group of the lipid. Overall, these models suggest that Sfh3 may recognize PI4P through a polar interaction network similar to that identified in the Sfh3-PI complex or via a different modality. In both cases, the R107 residue is involved in this interaction, consistent with our experimental data showing its critical role in PI4P transfer.

### Sfh3 efficiently exchanges lanosterol for PI

Our assays indicated that Sfh3’s ability to deliver DHE to acceptor membranes is inhibited by PI, lanosterol, and zymosterol but activated by PI4P present in these membranes **(Figs. 1 and 2)**. We also noted that PI4P activation of DHE transfer by Sfh3 is inhibited by PI (**Fig. 4)**. Moreover, our structure-function study confirmed that the DHE-transfer activity is inhibited by PI due to its recognition by Sfh3, and this protein recognizes PI4P weakly compared to PI **(Fig.5)**. Collectively, these data suggested that Sfh3 has a higher affinity for PI and ergosterol intermediates than for DHE and PI4P. To test this further, we measured the extent to which PI, PI4P, DHE, lanosterol, and zymosterol provide thermal stability to Sfh3 using nano differential scanning fluorimetry. Empty Sfh3 or in complex with a ligand was subjected to increased temperature and the denaturation of Sfh3 was followed by measuring its intrinsic fluorescence at 330 and 350 nm; determining the first derivative of the F_350_/F_330_ ratio with respect to temperature allowing identifying inflection point (IP) that corresponds roughly to the melting temperature (T_m_) at which half of the protein was unfolded **(Fig.5I)**. No or low increase in melting temperatures (ΔT_m_) were observed when Sfh3 was loaded with DHE and PI4P (0 ± 0.1°C and 0.8 ± 0.3 °C, respectively), compared to the value obtained when Sfh3 was empty **(Fig.5J)**, indicating that these ligands poorly stabilized the protein. In contrast, higher increases were obtained with Sfh3 in complex with PI (ΔT_m_ = 5.5 ± 0.03 °C), zymosterol (ΔT_m_ = 7.4 ± 0.1 °C), and lanosterol (ΔT_m_ = 9.7 ± 0.1 °C). We concluded that the relative affinity of Sfh3 for its ligands follows this order: DHE < PI4P < PI < zymosterol < lanosterol. In yeast cells, where all these lipid ligands are present, this implies that Sfh3 might preferentially transport ergosterol biosynthesis intermediates rather than ergosterol and may exchange these sterols efficiently with PI.

To investigate this, we devised an assay to measure Sfh3’s ability to transfer DHE, lanosterol, and PI individually or under exchange conditions **(Fig.6a)**. Sfh3 was mixed for 20 minutes with a 1:1 mixture of “heavy” L_A_ liposomes (loaded with sucrose, containing or lacking 5% PI, and doped with 2% NBD-PE) and “light” L_B_ liposomes (loaded with HK buffer, containing or lacking 5% sterol and doped with 2% Rhod-PE, see the Materials and Methods section for more details about preincubation steps). Controls were performed without Sfh3. After ultracentrifugation on a sucrose gradient, we collected bottom and top fractions, measured the fluorescence of NBD-PE and Rhod-PE in each fraction to estimate the cross-contamination between L_A_ and L_B_ liposomes, and then analyzed the lipid content of these fractions by LC-MS/MS following lipid extraction. For all conditions tested and analyzed below, fluorescence measurements indicated low cross-contamination, as 95% of L_A_ and L_B_ liposomes were recovered in the bottom and top fractions (e.g., **Fig. S8A, B**), and LC-MS/MS systematically confirmed this (**Fig. 6B)**.

**Figure 6.**
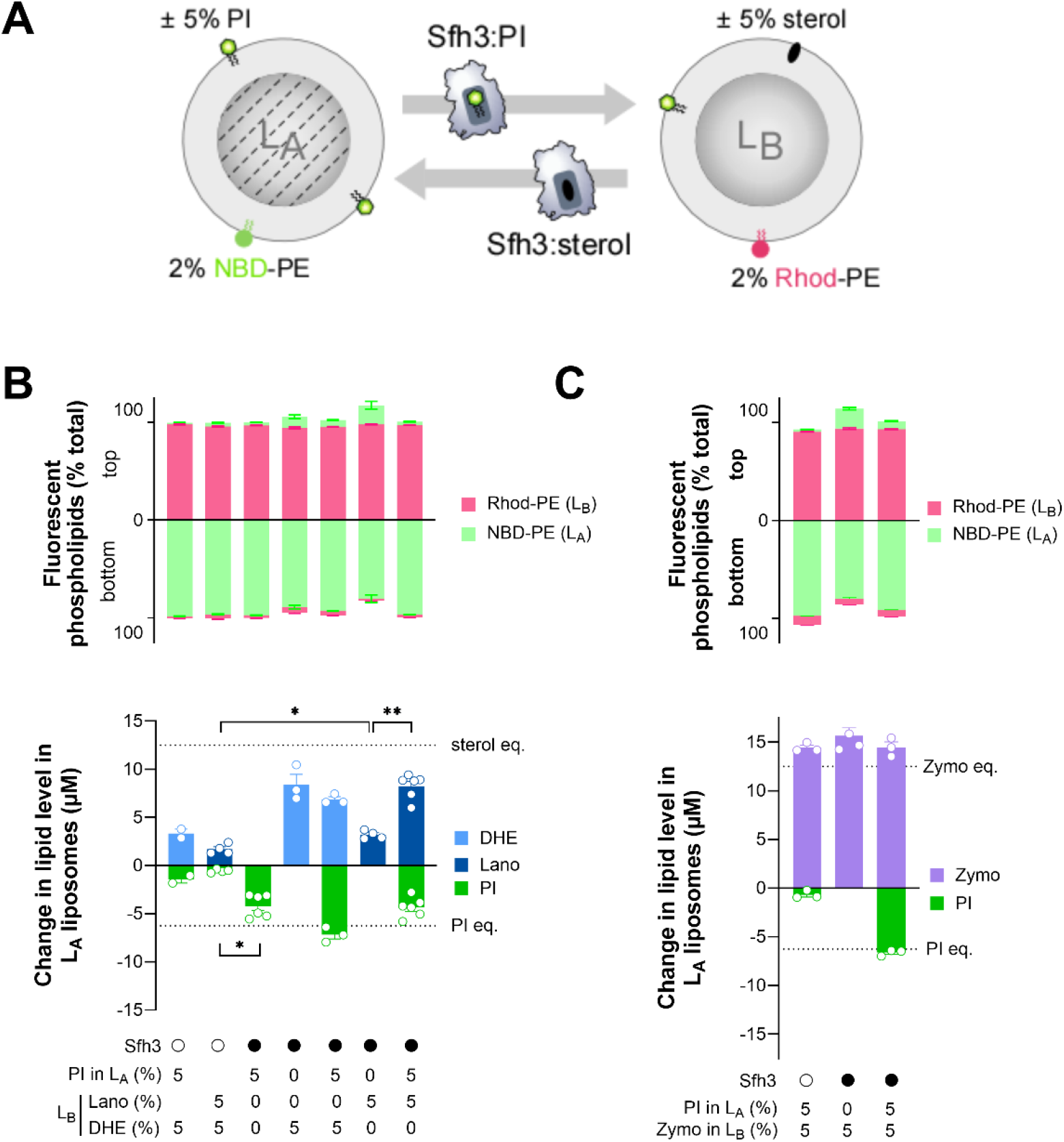
Sfh3 exchanges lanosterol and PI between membranes. **(A)** Principle of the LC-MS/MS-based lipid transfer assay. “Heavy” sucrose-loaded L_A_ liposomes (500 µM lipids) labeled with 2% NBD-PE, containing or not 5% PI, were incubated with the same amount of “light” sucrose-free L_B_ liposomes, labeled with 2% Rhod-PE, containing or not 5% sterol, and Sfh3 (500 nM). As a negative control, L_A_ and L_B_ liposomes containing 5% PI and 5% sterol, respectively, were mixed in the absence of Sfh3. After centrifugation on a sucrose gradient, the bottom and top fractions were collected, lipids were extracted from each fraction, and the lipids were quantified by LC-MS/MS. **(B)** Transfer experiments with L_A_ liposomes containing or not 5% PI and L_B_ liposomes containing or not 5% DHE or lanosterol. The top panel shows the relative amount of L_A_ and L_B_ liposomes in B+M1 (bottom) and T+M2 (top) fractions (in percentage) determined by quantifying NBD-PE and Rhod-PE using LC-MS/MS. The bottom panel shows the gain of sterol and loss of PI in L_A_ liposomes (in µM). Dashed lines indicate the theoretical gain and loss corresponding to equilibration of sterol and PI between L_A_ and L_B_ liposomes. Data are represented as mean ± s.e.m. (n = 4–6). Unpaired Mann– Whitney U test (\**p* < 0.05 and \*\**p* < 0.01). **(C)** Transfer experiments were repeated with L_A_ liposomes containing or not 5% PI and L_B_ liposomes containing 5% zymosterol in the absence or presence of Sfh3. NBD-PE, Rhod-PE, PI, lanosterol, and DHE amounts were quantified by LC-MS/MS in bottom and top fractions to assess how L_A_ and L_B_ liposomes were separated and how much PI and zymosterol were transferred by Sfh3. Data are represented as mean ± s.e.m. (n = 3).

First, we observed that without Sfh3, using L_A_ and L_B_ liposomes containing PI and sterol, respectively, there was a spontaneous transfer of 3.3 µM DHE and 1.7 µM lanosterol from L_B_ to L_A_ liposomes—representing 13% and 7% of their initial pools in LB membrane—along with a minor transfer of PI (0.5–1.5 µM, 2–6%) in the opposite direction. Subsequently, we found that Sfh3, in non-exchange conditions, could transfer approximately 4.2 µM PI from L_A_ to L_B_ liposomes lacking sterol. This transfer was notable, given that Sfh3 had access only to PI in the outer leaflet of L_A_ liposomes and that a maximum of 6.25 µM PI can be equilibrated between the outer leaflets of L_A_ and L_B_. Next, we tested Sfh3’s ability to transfer DHE from L_B_ to L_A_ liposomes, with or without PI. Sfh3 transferred 8.4 µM DHE to PI-free L_A_ liposomes, which was 2.5 times the spontaneous transfer. This value was close to the value expected if DHE was completely equilibrated between liposomes (*i.e.*, 12.5 µM, assuming that the entire DHE pool is accessible to Sfh3 as it rapidly flip-flops across the membrane ^38^). Under exchange conditions, with L_A_ liposomes containing PI, Sfh3’s capacity to transfer DHE was reduced by 30%; in contrast, its PI transfer activity was significantly enhanced by DHE. These data, aligning with kinetics measurements showing that PI inhibits Sfh3’s capacity to transfer DHE (Fig. 1) and DHE boosts its PI transfer activity (Fig. 3), confirm the validity of our experimental approach.

Once we established the protocol, we tested how Sfh3 transferred lanosterol under non-exchange and exchange conditions with PI. We observed that Sfh3 transferred 3.2 µM of lanosterol from L_B_ liposomes to PI-free L_A_ liposomes. This value corresponded to less than twice the quantity of spontaneously transferred lanosterol and was far from the value expected if lanosterol was completely equilibrated between L_A_ and L_B_ liposomes (considering that lanosterol also flip-flops^39^ and is fully accessible to Sfh3). This suggested that Sfh3 has a low basal lanosterol transfer activity. In contrast, when PI and lanosterol were present in L_A_ and L_B_ liposomes, respectively, Sfh3 transferred 4.4 times more lanosterol while its PI transfer activity was slightly increased **(Fig. 6B)**. We conclude that Sfh3 can individually transfer PI and lanosterol between membranes but transfers lanosterol much more efficiently in the presence of PI, likely via a lanosterol/PI exchange.

To complement our analysis, we examined whether Sfh3 could exchange PI and zymosterol between L_A_ and L_B_ liposomes doped with 5% PI and 5% zymosterol, respectively. With the protein present, we observed efficient PI transfer from L_A_ to L_B_ liposomes and substantial zymosterol transfer in the reverse direction, resulting in complete equilibration of the two lipid ligands between membranes **(Fig. 6C)**. However, even without PI or Sfh3, zymosterol was fully equilibrated. Because liposomes were properly separated in these assays and no spontaneous PI transfer occurred without the protein, we concluded that zymosterol is intrinsically prone to rapidly move spontaneously between membranes. Therefore, we cannot conclude that Sfh3 exchanges zymosterol with PI in addition to lanosterol.

### Sfh3 performs lanosterol/PI exchange under conditions mimicking azole-treated yeast

In yeast treated with a sub-lethal concentration of miconazole, half of the ergosterol is replaced by lanosterol in membranes ^16^ ^18^. We reasoned that analyzing Sfh3 transfer activity *in vitro* in a system that recapitulates these abnormal sterol conditions might help explain why Sfh3 provides resistance to azoles. Using our PI 4-kinase-based assays, we first analyzed how Sfh3 transfers PI between L_A_ and L_B_ liposomes containing 5% DHE and 5% lanosterol to mimic the sterol content of membranes of azole-treated yeast. We found that Sfh3 activity was similar to that measured with liposomes containing 10% DHE **(Fig. 7A)**. In contrast, Sfh3 could not transfer PI4P between membranes containing both DHE and lanosterol **(Fig. 7B)**, whereas Osh4’s activity remained unaffected. These data suggest that Sfh3 can transfer PI but not PI4P in azole-treated yeast.

**Figure 7.**
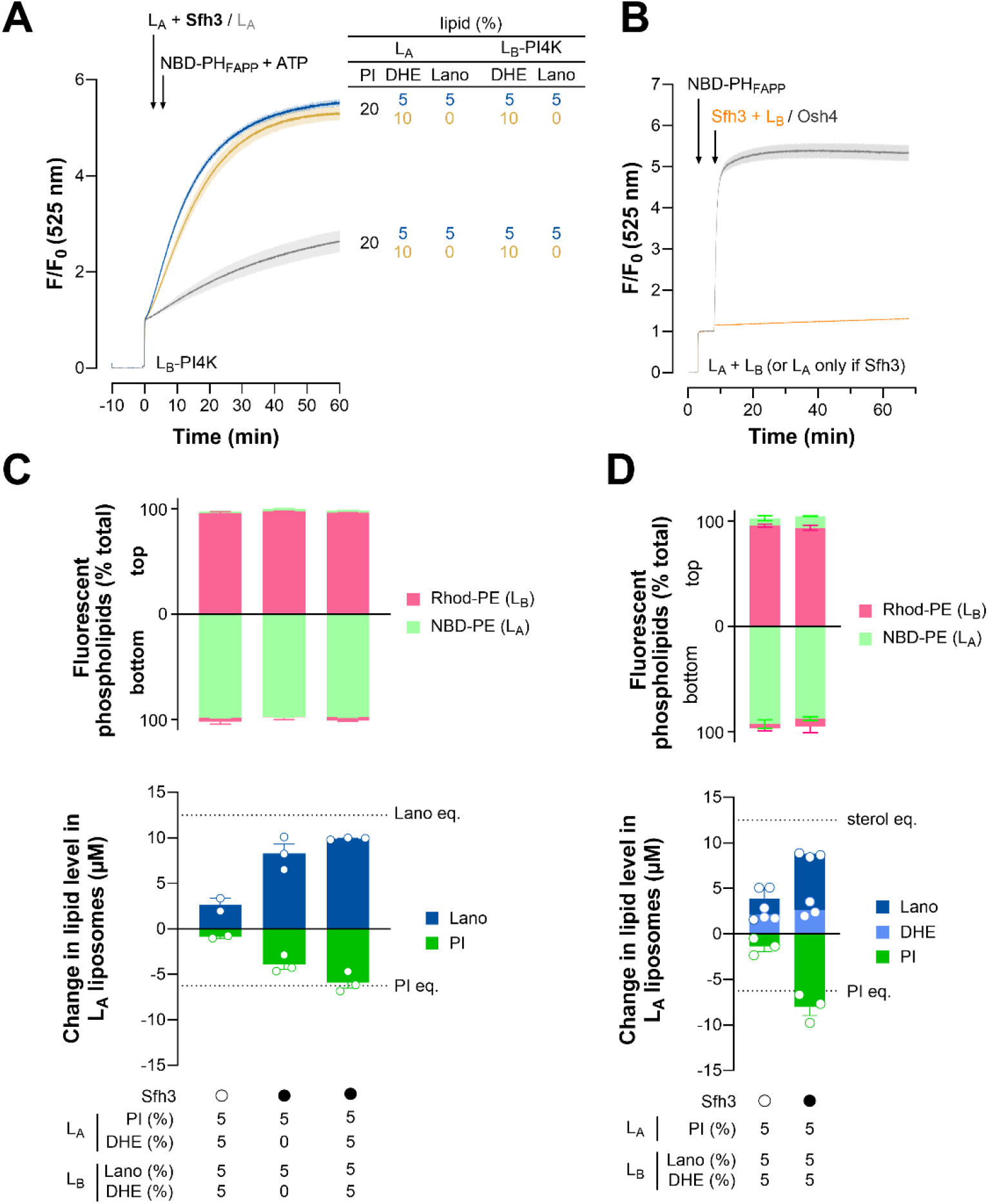
Sfh3 acts as a PI/lanosterol exchanger in conditions mimicking azole-treated cells. **(A)** PI-transfer assay. L_A_ liposomes composed of DOPC/PI/DHE/Rhod-PE (68:20:10:2) were pre-incubated, or not, with Sfh3 for 1 h at 30 °C and then added to L_B_-PI4K liposomes, composed of DOPC/DHE/MPB-PE (87:10:3), and coated with PI-4 kinase (P/L = 1/400). Alternatively, L_A_ and L_B_ liposomes, containing 5% DHE and 5% lanosterol instead of 10% DHE, were tested. Thirty seconds later, NBD-PH_FAPP_ (250 nM) and ATP (100 µM) were injected together. Final concentrations: Sfh3, 200 nM; L_A_ and L_B_ liposomes, 200 µM lipids each. Fluorescence was measured at 525 nm (λ_ex_ = 460 nm) at 30 °C with constant stirring and normalized to the signal (F_0_) measured immediately after the last injection. Each trace is the mean ± s.e.m. of several kinetic traces (n =3). **(B)** PI4P-transfer assay. L_A_ liposomes composed of DOPC/PI4P/DHE/lanosterol/Rhod-PE (83:5:5:5:2) were pre-incubated with Sfh3 for 1 h at 30 °C, then added to L_B_ liposomes, composed of DOPC/DHE/lanosterol (90:5:5), in the presence of NBD-PH_FAPP_. Osh4 was directly added to the L_A_ and L_B_ liposomes, mixed with NBD-PH_FAPP_. Each trace is the mean ± s.e.m. of several kinetics (n = 4). **(C)** LC-MS/MS lipid transfer assay. L_A_ and L_B_ liposomes (500 µM lipid each), respectively doped with 5% PI and 5% lanosterol, containing or not 5% DHE, were mixed with Sfh3 (500 nM). Negative controls used L_A_ and L_B_ liposomes with 5% PI and lanosterol plus 5% DHE, without Sfh3. After centrifugation on a sucrose gradient, fractions were collected, lipids were extracted, and the lipids were quantified. The amounts of L_A_ and L_B_ liposomes in each fraction, determined by NBD-PE and Rhod-PE quantification, are shown in the top panel. Lanosterol, DHE, and PI gain/loss in L_A_ liposomes were reported in the bottom panel. Data are mean ± s.e.m. (n=2–3). **(D)** The experiment described in (C) was repeated with L_A_ liposomes doped with 5% PI and L_B_ liposomes containing 5% DHE and 5% lanosterol in the absence or presence of Sfh3. The amounts of NBD-PE, Rhod-PE, PI, lanosterol, and DHE were quantified in bottom and top fractions to assess how the L_A_ and L_B_ liposomes were separated and how much PI, lanosterol, and DHE were transferred by Sfh3. Data are represented as mean ± s.e.m. (n = 3).

Because our data suggested that Sfh3 might act preferentially as a lanosterol/PI exchanger in cells, we tested whether Sfh3 can do so in the presence of ergosterol. We analyzed how Sfh3 performs PI/lanosterol exchange between L_A_ and L_B_ liposomes, respectively containing 5% PI and 5% lanosterol, and doped or not with 5% DHE. In the absence of DHE, we found that Sfh3 transfers 20% of the PI initially contained in L_A_ liposomes and 30% initially contained in L_B_ liposomes along opposite routes, confirming that it acts as a PI/lanosterol exchanger **(Fig. 7C)**. In the presence of DHE, the PI/lanosterol exchange activity of Sfh3 was not affected and was even slightly higher, indicating that the presence of DHE does not impact this process. To confirm these data, we examined whether Sfh3 could selectively transfer lanosterol from L_B_ liposomes containing both 5% lanosterol and 5% DHE in exchange for PI. We found that this was the case, as Sfh3-mediated lanosterol transfer from L_B_ to L_A_ liposomes was higher than spontaneous lanosterol transfer, whereas Sfh3-mediated DHE transfer was similar to spontaneous DHE transfer under conditions where Sfh3 transferred PI in the opposite direction **(Fig. 7D)**. These data suggest that Sfh3 can preferentially exchange lanosterol and PI in miconazole-treated yeast.

### The ability of Sfh3 to confer azole resistance involves its PI/lanosterol exchange activity

The Sfh3^E235A/K267A^ mutant behaves markedly differently from Sfh3 WT across various cellular contexts and, notably, does not confer resistance to azole treatment in yeast^16^, possibly because it fails to recognize PI. To test this, we examined how mutations that either affect or do not affect Sfh3’s ability to transfer PI in our *in vitro* assay alter its ability to protect yeast against miconazole. To do this, we introduced these mutations, including the E235A/K267A double mutation, into the chromosomal copy of SFH3, which was expressed as an mScarlet fusion. The resulting yeast strains, as well as a *sfh3Δ* strain, were then challenged with 50 ng/mL miconazole for four days in the dark. The experiment confirmed that the *sfh3Δ* strain is more susceptible to miconazole than wild-type cells. We also confirmed that the E235A/K267A double mutation disrupts Sfh3’s ability to protect yeast against miconazole. We observed a similar effect with the K98A, L181Y, T264W, and Q230A/T264W mutations. In contrast, expression of Sfh3 bearing the L158Y, Q230A, or E235A mutations did not induce this defect. Because the first category of mutants—except Sfh3^K98A^—was strongly defective in PI transfer, whereas the second category was not, we conclude that Sfh3’s ability to confer resistance to miconazole is linked to its ability to recognize PI.

To better understand why the capacity of Sfh3 to provide azole resistance is linked to its capacity to recognize PI, we further characterized the Sfh3^E235A/K267A^ mutant. We noted that this mutant transported DHE faster than Sfh3 WT in the absence and presence of PI. To strengthen these results, we first measured its ability to transport DHE from L_A_ to L_B_ liposomes containing increasing concentrations of PI, and performed similar measurements with Sfh3 WT. We found that Sfh3^E235A/K267A^ transported DHE at PI concentrations from 0% to 20% much faster than Sfh3 WT, although its activity decreased slightly at 20% PI **(Fig.8B, Fig. S4B)**. This result confirms that this mutant does not recognize PI as a competitive ligand and can transfer DHE at high PI concentrations, suggesting that it can fully transport ergosterol in yeast. Next, we examined the ability of Sfh3^E235A/K267A^ to transport DHE from L_A_ liposomes to L_B_ liposomes containing either 0% or 5% lanosterol **(Fig. 8C)** and found that lanosterol blocked its activity, indicating that it can properly recognize this lipid ligand. These data show that the Sfh3^E235A/K267A^ mutant’s sterol-binding capacity is not changed by the double mutation, and that its distinct behavior from Sfh3 WT in azole-treated yeast is not due to a sterol recognition defect.

**Figure 8.**
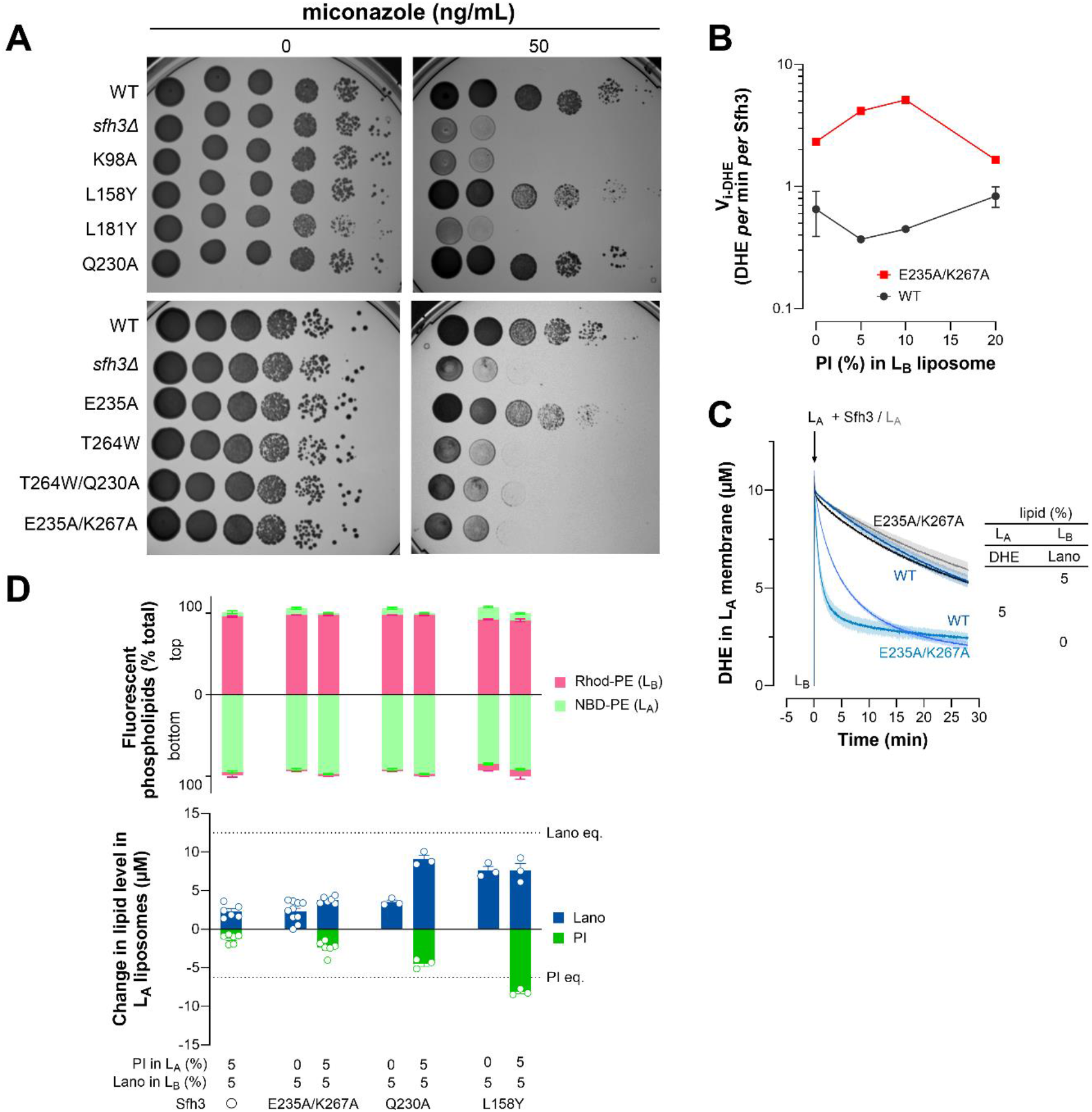
Sfh3^E235A/K267A^ mutant is defective in operating lanosterol/PI exchange. **(A)** Test of sensitivity of yeast strains expressing Sfh3 variants to miconazole. Ten-fold serial dilutions of indicated strains were spotted onto YPD plates supplemented or not with 50 ng/mL miconazole. Plates without miconazole were incubated at 37 °C in the dark for 2 days, while plates containing miconazole were incubated for 4 days. All strains except *sfh3Δ* express either WT or mutant versions of Sfh3 from the endogenous chromosomal locus, as indicated, tagged C-terminally with mScarlet. **(B)** Initial DHE transfer rate measured for Sfh3 and Sfh3^E235A/K267A^ (1.5 µM) between L_A_ liposomes (200 µM) composed of DOPC/DHE/DNS-PE (92.5:5:2.5) and L_B_ liposomes (600 µM) made of DOPC and containing no PI or 5, 10, or 20% PI. **(C)** Sfh3 or Sfh3^E235A/K267A^ was pre-incubated with L_A_ liposomes composed of DOPC/DHE/DNS-PE (92.5:5:2.5, 200 µM total lipids) for 1 h, and the mixture was injected into a cuvette containing L_B_ liposomes made of DOPC, containing or not 5% lanosterol. Final concentrations: L_A_ liposomes, 200 µM; L_B_ liposomes, 600 µM; Sfh3, 1.5 µM. Each trace is the mean ± s.e.m. of several kinetics (n = 2–4). **(D)** LC-MS/MS-based lipid transfer assay. The experiment was performed using L_A_ liposomes with or without 5% PI and L_B_ liposomes doped with 5% lanosterol. The liposomes were mixed with Sfh3^E235A/K267A^, Sfh3^Q230A^, or Sfh3^L158Y^ mutant. The relative amounts of L_A_ and L_B_ liposomes in the bottom and top fractions are shown in the top panel. The bottom panel shows the gain of lanosterol and loss of PI in LA liposomes. Data are presented as mean ± s.e.m. (n = 3–4).

Next, because Sfh3 can mediate lanosterol/PI exchange but the E235A/K267A double mutation prevents PI transfer, we asked whether this mutation affects the protein’s lanosterol/PI exchange activity. Using our LC-MS/MS approach, we found that Sfh3^E235A/K267A^ could slightly transport lanosterol from L_B_ liposomes to PI-free L_A_ liposomes, as observed for Sfh3 WT. We repeated the experiment with L_A_ liposomes containing 5% PI and found that this mutant weakly transported PI from L_A_ to L_B_ liposomes, as expected, and did not substantially transfer lanosterol in the opposite direction, contrary to Sfh3 WT **(Fig. 6B).** We also tested the Sfh3^Q230A^ and Sfh3^L158Y^ mutants, which protect yeast against miconazole. We found that Sfh3^Q230A^ had low lanosterol transfer activity in the absence of PI but was strongly activated by PI, similar to Sfh3 WT. The Sfh3^L158Y^ mutant behaved slightly differently: it showed high basal lanosterol transfer activity even without PI, possibly due to lower sterol affinity, as suggested by our structure-function analysis. Nevertheless, it was still able to efficiently exchange lanosterol for PI **(Fig. 8D)**. Collectively, these results suggest that Sfh3 becomes defective in conferring azole resistance in yeast if it cannot mediate lanosterol/PI exchange.

## Discussion

Here, we have dissected the lipid-transfer properties of Sfh3 to gain insight into its cellular role. After establishing a reliable assay to measure Sfh3’s ability, in a monomeric state, to transfer DHE, a natural ergosterol analog, we tested whether it could act as an ergosterol/PI exchanger. We found that its capacity to transfer DHE is not activated but inhibited by PI under exchange conditions in which these lipids are present in small amounts in separate membranes. Under similar conditions, we found that Osh4 transfers DHE up to 20-fold faster once its second ligand, PI4P, is present, as previously shown^31^. We next found that PI inhibits Sfh3’s capacity to transfer DHE because it is specifically recognized by the protein, not because it provides negative charges to membranes; indeed, substituting it with another anionic lipid, PS, stimulates DHE transfer. In fact, Sfh3-mediated DHE transfer can be restored when the acceptor membranes contain a high quantity of PI (20%), comparable to that of yeast endomembranes, suggesting that the negative charge provided by PI to membranes counterbalances its inhibitory action. Eventually, analysis of Sfh3 mutants showed that the inhibitory effect of PI on the DHE transfer activity of Sfh3 correlates with the protein’s capacity to transfer PI. We conclude that the ergosterol transfer activity of Sfh3 is not activated by PI inside yeast.

Because Sfh3 can recognize lanosterol and zymosterol^21^, we compare this capacity to its ability to recognize ergosterol. Under exchange conditions, we found that lanosterol and zymosterol, but not ergosterol, block the DHE transfer activity of Sfh3 even when they are 10-fold less abundant than DHE. By comparison, we observed no inhibition of STARD3-mediated DHE transfer activity. We also noted that the PI4P transfer activity of Osh4, activated by DHE, remains insensitive to lanosterol. It is unclear how Sfh3 could recognize lanosterol and ergosterol differently^21^, but our data suggest that, compared to STARD3 and Osh4, Sfh3 is structurally poised to recognize ergosterol intermediates with the highest affinity. This suggested it might preferentially transfer these sterols within yeast, even if they are present in minor amounts^40^.

Next, we measured Sfh3’s capacity to transfer PI from a first to a second liposome population by quantifying PI-to-PI4P conversion in the second population. In the absence of DHE, the PI transfer activity of Sfh3 was lower than that of Sec14, as observed previously in assays likely performed with sterol-free membranes^15, 41^. In the presence of DHE, we found that Sfh3’s activity nearly matched Sec14’s, likely because it can exchange PI for its second ligand, as Sec14 does with PC^42–43^. Sfh3 delivers PI even more effectively to membranes containing lanosterol, possibly due to its higher affinity for this sterol. However, we found that Sfh3 can transfer PI4P between membranes, and lanosterol (but not DHE or zymosterol) inhibits this process, complicating our analysis based on PI4P quantification in a single liposome population. Still, we can conclude that Sfh3’s PI transfer activity *per se* is facilitated by sterol.

We further investigated whether Sfh3 can efficiently transfer PI4P and concluded that it may not be a robust PI4P transporter in cells, unlike ORPs^1, 8^. Compared to Osh4, Sfh3 exhibits a modest PI4P transfer activity *in vitro* that is strongly attenuated when membranes are enriched with PI. Additionally, we observed that PI4P, unlike PI, activates the DHE-transfer activity of Sfh3, but that this effect is inhibited by PI. Moreover, mutations in Sfh3 (E235A, K98A, R107A, L158Y) that have minimal effects on its PI transfer activity substantially decrease or abolish its PI4P transfer activity. All these data suggest that Sfh3 interacts more weakly with PI4P than with PI. Presumably, Sfh3 recognizes the PI moiety of PI4P through weaker interactions than those involved in PI recognition, and this is not compensated by additional contacts between Sfh3 and the phosphate group at the 4-position of the inositol ring. Mutating R107 specifically abolishes Sfh3’s PI4P transfer activity. Because this residue is not conserved in Sec14, which lacks PI4P transfer activity, it suggests that R107 enables Sfh3 to recognize PI4P. Interestingly, Sec14-like protein 2 (SEC14L2) has recently been shown to transport PI4P and PI3P in cells^44^ despite its strong capacity to bind to PI^45–46^, suggesting it has the highest affinity for phosphoinositides, unlike Sfh3. Further structural and functional comparisons between Sec14, Sfh proteins^12^, and Sec14-like proteins (SEC14L2 but also SEC14L6)^47^ may help determine how well these proteins can recognize PI and phosphoinositides.

Our intuition about the relative affinity of Sfh3 for its lipid ligand was confirmed by nanoDSF, showing that Sfh3 has a preference for lanosterol, zymosterol, and PI rather than for DHE and PI4P. In yeast, where all these lipid species are present, this suggests that Sfh3 preferentially recognizes PI and ergosterol intermediates and exchanges them. Supporting this assumption, we established that Sfh3 is a lanosterol/PI exchanger by showing that, unlike DHE, Sfh3 delivers lanosterol more efficiently to an acceptor membrane when PI is present and transfers PI back to the membrane that initially contained lanosterol. However, we could not prove that Sfh3 also acts as a zymosterol/PI exchanger because of high spontaneous zymosterol transfer in our assay. Importantly, we found that Sfh3 can mediate PI/lanosterol exchange in the presence of DHE without displacing DHE. We conclude that Sfh3 can exchange lanosterol for PI in yeast, despite the prominence of ergosterol.

The finding that Sfh3 can transfer ergosterol, lanosterol, PI, and PI4P with varying efficiencies suggests that it might function under different regimes in yeast. One possibility is that its dual PI/PI4P transport capacity enables it to regulate specific cellular PI4P pools in a sterol-dependent manner. In our experimental setup—designed to measure PI transfer and mimicking an interface between the ER and a compartment equipped with PI4-kinase (e.g., the Golgi apparatus, vacuole, or plasma membrane)—we observed that changes in sterol levels and types modulate PI4P accumulation in that compartment by tuning Sfh3’s ability to deliver PI and remove PI4P. We therefore propose that, in yeast, Sfh3 may weakly deliver PI into, while substantially extracting PI4P from, compartments with low ergosterol or ergosterol intermediate levels, thereby lowering PI4P signal, and does the opposite when ergosterol or ergosterol intermediate levels increase **(Fig. 9A)**. This model might explain enigmatic observations obtained by Bankaitis team, showing that Sfh3 can both promote PI4P synthesis in yeast and antagonize Sec14/Pik1-dependent PI4P signaling at the Golgi ^15^. The notion that Sfh3’s function is to regulate PI4P levels was proposed by the same group to explain its capacity to inhibit the mobilization of neutral lipids and sterols from LDs during yeast sporulation ^15^. However, it is unclear whether this occurs in a sterol-dependent manner because much of the study relies on analysis of the Sfh3^T264W^ mutant, which is not exclusively unable to recognize PI as reported before^15^, but also sterol and PI4P, as we show here. Our model also suggests that Sfh3 can regulate PI4P signaling while, in turn, moving sterol. This might explain why this protein is localized in vCLIP^13–14^ in the context of microlipophagy. Because the formation of microdomains rich in ergosterol^48–49^ and PI4P^50^ in the vacuolar membrane is needed for this process, we propose that Sfh3 fine-tunes the levels of two lipids. Sfh3 might also, or alternatively, limit the accumulation of ergosterol intermediates in that membrane, which has been shown to impair sterol-rich domain formation^51^. Further investigations are needed to explore these possibilities.

**Figure 9.**
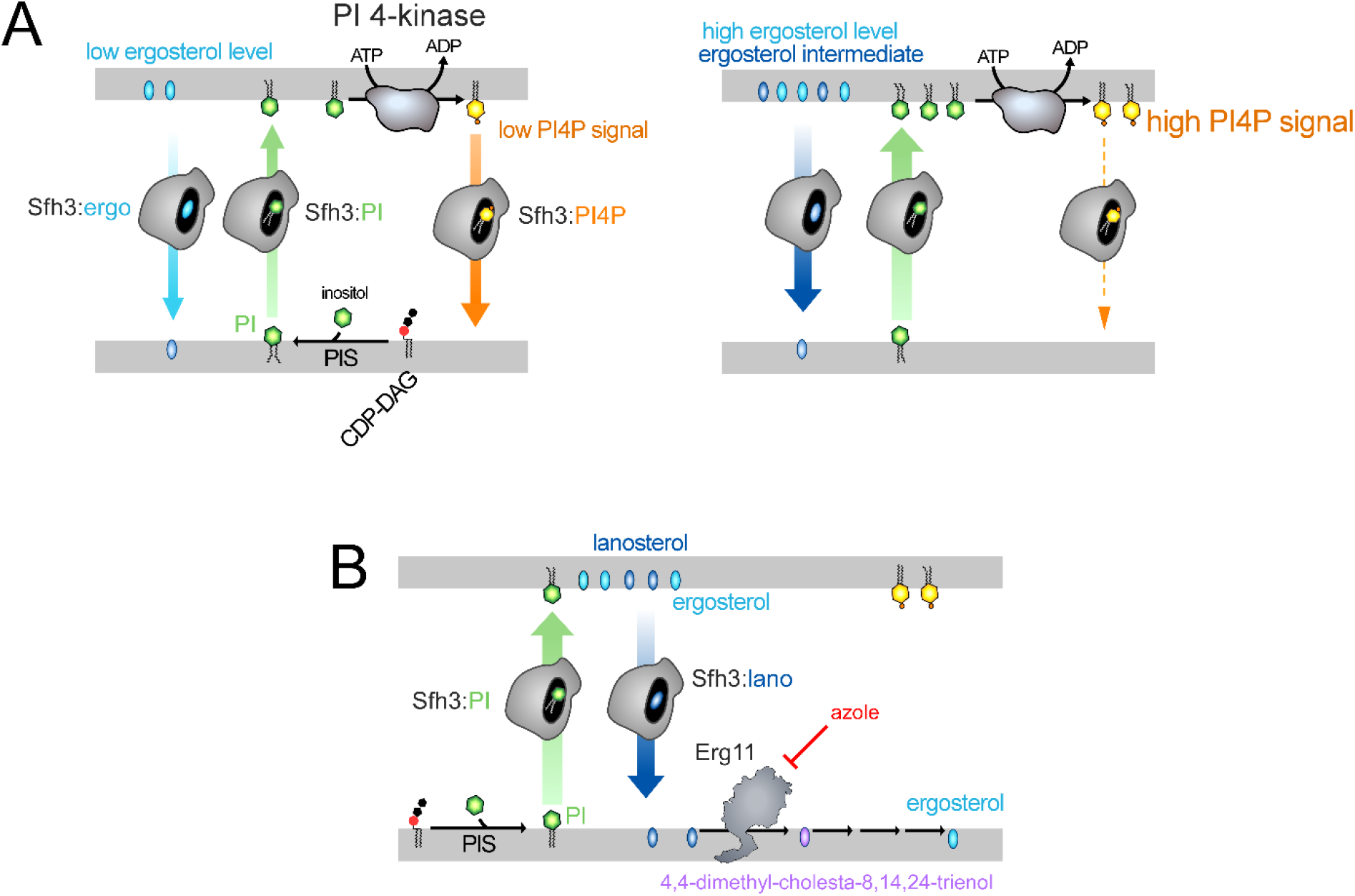
Model on the potential cellular role(s) of Sfh3. **(A)** Sfh3 could regulate PI4P levels in a sterol-dependent manner. Sfh3 might inefficiently deliver PI to a membrane equipped with PI 4-kinase activity if that membrane is poor in sterol, because sterol/PI exchange is limited, while removing PI4P efficiently from this membrane once it is synthesized. This would result in low PI4P levels. In the presence of higher ergosterol concentrations (or ergosterol intermediates such as lanosterol), Sfh3 could more efficiently supply PI via PI/sterol exchange, allowing higher PI4P production while extracting PI4P less efficiently. This would increase PI4P levels. **(B)** Potential role of the Sfh3 protein during azole treatment. In response to partial inhibition of Erg11 by azoles, which leads to lanosterol accumulation and spreading in different membranes, Sfh3 could extract lanosterol from these membranes in exchange for PI. Lanosterol would thus be delivered to the ER, maintaining a high substrate level for Erg11 and counteracting azole inhibition. The continuous synthesis of PI from CDP-DAG and inositol would drive continuous PI/lanosterol exchange cycles. Lanosterol would then be continuously reintegrated into the ergosterol biosynthesis pathway, and the yeast cells would be detoxified, becoming resistant to azole treatment.

Given that Sfh3 can preferentially recognize lanosterol and PI, we also propose that in miconazole-treated yeasts, Sfh3 engages in an exclusive regime of lanosterol/PI exchange as intracellular lanosterol concentration rises, conferring resistance to this drug. This model is further supported by our analysis of the Sfh3^E235A/K267A^ mutant, previously studied to understand Sfh3’s role in azole resistance^16, 21^. We confirm that it cannot transfer PI and show that it is not defective in transporting sterol, clarifying contrasting data^11, 16^. Instead, we observe that Sfh3^E235A/K267A^ transfers DHE more efficiently than Sfh3 WT in the presence and even in the absence of PI (for reasons that might be linked to sterol or PC recognition, as E235A and K267 residues might interact with sterol or with the 1-phosphate group of PC, considering molecular docking and structural analysis^11, 21^ and the structure of Sfh3:PI complex^20^). Most importantly, we reveal that this mutant recognizes lanosterol but cannot transfer it rapidly under exchange conditions with PI. Given this observation and that Sfh3, but not Sfh3^E235A/K267A^, provides yeast with resistance to sublethal azole doses while reducing lanosterol levels^16^, we propose the following model. Sfh3 might exchange PI, synthesized in the ER, with lanosterol spread in other compartments^40^ and deliver lanosterol to the ER. This exchange might maintain high lanosterol levels for Erg11, counteracting azole inhibition and allowing more lanosterol to eventually enter the ergosterol biosynthesis pathway. This Sfh3-mediated process could thus allow yeast detoxification and survival in the presence of azole **(Fig. 9B)**.

In conclusion, our study provides new insights into Sfh3, suggesting that it might preferentially exchange sterol intermediates for PI intracellularly, possibly to regulate PI4P-dependent processes and ergosterol biosynthesis; this mechanism might explain why Sfh3 confers azole resistance in yeast. However, our study cannot clarify whether Sfh3 provides the *Δerg6* strain with resistance to aureobasidin A^28–29^ by transferring zymosterol between membranes in a PI-dependent manner. More broadly, our study shows that an LTP can recognize and transfer multiple lipid ligands with varying efficiencies and suggests that its activity within cells likely depends on ligand concentrations in the cellular environment. Therefore, our findings indicate that merely identifying the lipid ligands of LTPs is insufficient. Confirming previous conclusions on ORPs^30^, we found that it is critical to determine how LTPs recognize their lipid ligands hierarchically and to test mechanistic scenarios using membranes with complex lipid compositions.

## Material and methods

### Protein expression and purification from *E. coli*

GST-Sfh3 and mutants were expressed in *E. coli* (BL21-GOLD(DE3)) competent cells (Stratagene) grown in Luria-Bertani broth (LB) at 30 °C overnight after induction with 1 mM isopropyl β-D-1-thiogalactopyranoside (IPTG). When the optical density of the bacterial suspension, measured at 600 nm, reached 0.6—0.7, bacterial cells were harvested and resuspended in cold TN buffer (50 mM Tris-HCl, pH 7.4, 150 mM NaCl, 1 mM DTT) supplemented with 1 mM PMSF, 10 µM bestatin, 1.6 µM pepstatin A, and cOmplete® EDTA-free protease inhibitor tablets (Roche). Cells were lysed in a Cell Disruptor TS SERIES (Constant Systems Ltd.), and the lysate was centrifuged at 186,000 × g for 1 h 30. The supernatant was then applied to Glutathione Sepharose 4B (Cytiva) for 3 h 30 at 4 °C. The beads were washed 4 times with TN buffer lacking protease inhibitors, then incubated overnight at 4 °C with thrombin to cleave Sfh3 from the GST domain. After several cycles of centrifugation and bead washing, the construct was recovered in the supernatant, concentrated, and injected onto an XK-16/70 column packed with Sephacryl S-200 HR for size-exclusion chromatography. Fractions containing ∼100% pure Sfh3 were pooled, concentrated, and supplemented with 10% (v/v) pure glycerol (Sigma). Aliquots were prepared, flash-frozen in liquid nitrogen, and stored at -80 °C. The protein concentration was determined by spectrophotometry at λ = 280 nm using an extinction coefficient ε = 52,495 M^-1^.cm^-1^. Sec14 and PI4K were purified as described in ^35^. cSTD3 was purified as described in ^52^. Osh4 and NBD-PH_FAPP_ were purified as described in ^31^.

### Lipids

18:1/18:1-PC (1,2-dioleoyl-*sn*-glycero-3-phosphocholine or DOPC), 16:0/18:2-PS (1-palmitoyl-2-linoleoyl-sn-glycero-3-phospho-L-serine or PLPS), soy PI (L-α-phosphatidylinositol), brain PI4P (L-α-phosphatidylinositol 4-phosphate), 18:1/18:1-PI4P, NBD-PE (1,2-dioleoyl-*sn*-glycero-3-phosphoethanolamine-N-(7-nitro-2-1,3-benzoxadiazol-4-yl)), Rhod-PE (1,2-dipalmitoyl-*sn*-glycero-3-phosphoethanolamine-N-(lissamine rhodamine B sulfonyl), 18:1 MPB-PE (1,2-dioleoyl-*sn*-glycero-3-phosphoethanolamine-N-[4-(p-maleimidophenyl)butyramide]), DNS-PE (1,2-dioleoyl-*sn*-glycero-3-phosphoethanolamine-N-(5-dimethylamino-1-naphthalenesulfonyl)), ergosterol, lanosterol and zymosterol were purchased from Avanti Polar Lipids. Yeast PI was obtained from Abitec (Larodan Research-Grade Lipids). Dehydroergosterol (DHE) was obtained from Sigma-Aldrich. The concentration of DHE in the stock solution in methanol was determined by UV spectroscopy using an extinction coefficient of 13,000 M^-1^.cm^-1^.

### Liposomes preparation

In glass tubes, lipids stored in CHCl_3_ or CHCl_3_/MeOH stock solutions were mixed at the desired molar ratio. The tubes were pre-warmed at 33 °C for 5 min; the solvent was then dried under a nitrogen stream for 25 min, and the tubes were placed in a vacuum chamber for 40 min to remove residual solvent. The lipid film was hydrated with 50 mM HEPES, pH 7.4, 120 mM K-Acetate (HK) buffer to obtain a multilamellar vesicle suspension. To prepare heavy liposomes, the lipid film was hydrated with 50 mM HEPES, pH 7.4, 220 mM sucrose buffer. The multilamellar vesicle suspensions were frozen and thawed 5 times, then stored at –20°C before being extruded on the day of the experiment through polycarbonate filters with 0.2 µm pore size using a mini-extruder (Avanti Polar Lipids) to obtain unilamellar vesicles (liposomes).

### Circular dichroism

The experiments were performed on a Jasco J-815 spectrometer at room temperature using a quartz cell with a path length of 0.05 cm (Starna Scientific Ltd.). Each protein was dialyzed in a Slide-A-Lyzer MINI dialysis device (MWCO 3.5 kDa) three times against 20 mM Tris, pH 7.4, 120 mM NaF buffer for 30 min to remove glycerol or DTT from the protein stock and to exchange the buffer. The samples were then ultracentrifuged at 100,000 × *g* for 20 min at 20 °C to pellet protein aggregates, and the supernatant was collected. Each CD spectrum represents the average of 10 scans recorded from λ = 190 to 260 nm, with a bandwidth of 1 nm, a step size of 0.5 nm, and a scan speed of 50 nm.min⁻¹. Protein concentration was determined at λ = 280 nm by spectrophotometry or densitometry after SDS-PAGE, using a BSA standard. A buffer control spectrum was subtracted from each protein spectrum. The percentages of secondary structure for each construct were estimated by analyzing their CD spectra (in the 190–250 nm range) using the BeStSel online method ^53^.

### Kinetics assays

All kinetics assays were carried out in a JASCO FP-8350 fluorimeter. The sample (volume 600 µL) was placed in a cylindrical quartz cell, continuously stirred with a small magnetic bar, and thermostated. At the indicated time, the sample was injected from stock solutions through a guide in the fluorimeter cover adapted for Hamilton syringes, to avoid interrupting the fluorescence recording.

### DHE transfer assay

Liposomes (termed L_A_, 4 mM lipids), composed of DOPC/DHE/DNS-PE (92.5:5:2.5, mol/mol), were pre-mixed with Sfh3 (30 µM) for 1 hour at 30°C under agitation. Thereafter, 30 µL of this mix was added to L_B_ liposomes (600 µM lipid, final concentration), either composed solely of DOPC or containing additional phospholipids (PI, PS, PI4P) or sterols (ergosterol, lanosterol, zymosterol), in 570 µL of HK buffer at 30 °C under agitation. In some cases, experiments were performed with an equal amount of L_A_ and L_B_ liposomes (200 µM) and 200 nM Sfh3. For DHE transfer assays performed with Osh4, 30 μl of L_A_ liposomes (200 μM total lipids, final concentration) were added to a suspension of L_B_ liposomes (200 μM or 600 μM total lipids) at 30 °C under continuous stirring in HK buffer (total volume 570 μL). After 2 minutes, 200 nM or 1.5 µM Osh4 was injected. DHE transport was monitored by measuring the dansyl signal at λ_em_ = 525 nm (bandwidth 5 nm) upon excitation at λ_ex_ = 310 nm (bandwidth 1 nm). At the end of the kinetic run, a final concentration of 7.5 mM methyl-β-cyclodextrin was added to fully extract DHE from the liposomes. The amount of DHE (in µM) transferred from L_A_ to L_B_ liposomes corresponds to 10 × ((F-F_0_)/(F_max_-F_0_)). F_max_ is the signal measured before adding Sfh3 or another LTP. DHE is quickly equilibrated between the inner and outer leaflets of liposomes (t_1/2_ of transbilayer movement <1 minute ^38^), implying that a pool of 10 µM DHE is accessible to Sfh3.

### PI transfer assay based on PI4P synthesis detection

On the day of the experiment, 100 µL of PI4K stock solution was applied to a Zeba column, equilibrated with freshly degassed HK buffer according to the manufacturer’s instructions, to remove DTT from the protein. The concentration of the eluted protein was determined by UV spectroscopy. DTT-free PI4K (2 µM final concentration) was then incubated with liposomes (L_B_, 800 µM lipids), of the desired composition and containing 3% MPB-PE, for 1 hour at 20 °C. The reaction was stopped by adding 1 mM DTT. Subsequently, a 150 µL aliquot was mixed with 420 µL of HK buffer supplemented with 10 mM MgCl₂ in a cuvette thermostated at 30°C. Nine minutes and 30 seconds later, NBD-PH_FAPP_ (250 nM) and 100 µM ATP were added simultaneously. Thirty seconds later, 30 µL of a suspension of liposomes, composed of DOPC/PI/Rhod-PE (78:20:2, mol/mol, L_A_ liposomes, 4 mM lipids) and pre-mixed with 200 nM Sec14 or Sfh3 for 1 hour, was injected. Fluorescence was measured at λₑₘ = 525 nm (bandwidth 10 nm) upon excitation at λₑₓ = 460 nm (bandwidth 1 nm) under constant stirring.

### PI4P transfer assay

A suspension (30 µL) of L_B_ liposomes (4 mM total lipid), composed of DOPC/PI4P/Rhod-PE (93:5:2, mol/mol), supplemented or not with PI or DHE, was diluted in 540 µL HK buffer. The mixture was placed in a cylindrical quartz cuvette, continuously stirred, and maintained at 30 °C. After 5 minutes, 250 nM NBD-PH_FAPP_ was injected. Five minutes later, a 30 µL suspension of L_A_ liposomes (4 mM lipids), composed solely of DOPC or containing additional lipid (DHE, lanosterol, zymosterol, or PI), incubated with Sfh3 (4 µM) for 1 hour at 30 °C under agitation, was injected. For experiments with Osh4 and Sec14, which do not require incubation, 200 nM of each protein was injected at 10 minutes into a mixture of L_A_ and L_B_ liposomes (200 µM lipid each) and NBD-PH_FAPP_ (250 µM), for a final HK volume of 600 µL. The NBD signal was measured at λ_em_ = 525 nm (bandwidth 10 nm) upon excitation at λ_ex_ = 460 nm (bandwidth 1 nm), with a time resolution of 0.5 s.

### LC-MS/MS-based transfer assays

This protocol is based on a differential flotation assay. “Heavy” liposomes (L_A_, 500 µM, final concentration) made of DOPC, doped with 2% NBD-PE and containing or not 5% PI, loaded with 50 mM HEPES, pH 7.4, 220 mM sucrose buffer, were mixed or not with Sfh3 (200 nM final concentration) for 1 h at 25 °C under an agitation of 1000 rpm and then added to “light” liposomes, composed of DOPC, and including 2% Rhod-PE and 0 or 5% sterol (L_B_, 500 µM) to initiate the transfer experiment. The sample (125 µL) was mixed for 20 minutes at 30 °C under constant agitation. Then, the sample was transferred to a polycarbonate tube adjusted to 7.5% sucrose by mixing 75 µL of a 20% (w/w) sucrose HK buffer, then overlaid with 200 µL of 5% (w/w) sucrose HK buffer, with 200 µL of a 2.5% (w/w) sucrose HK buffer, and finally with 200 µL of sucrose-free HK buffer. The suspension was centrifuged at 240,000 × *g* for 1 hour using an Optima Max XP ultracentrifuge with a TLS 55 swinging-bucket rotor (Beckman Coulter). The bottom (B, 200 µL), middle 1 (M1, 200 µL), middle 2 (M2, 200 µL), and top (T, 200 µL) fractions were collected. Each tube was imaged before and after ultracentrifugation to confirm that the heavy and light liposomes were, respectively, at the bottom and top of the sucrose gradient and were therefore well separated. The relative amounts of L_A_ and L_B_ liposomes in each fraction were then precisely quantified by mixing 10 µL of each fraction with 90 µL of HK buffer and measuring the fluorescence of NBD-PE and Rhod-PE. The bottom fraction was then pooled with the middle 1 fraction, and the top fraction was pooled with the middle 2 fractions. Lipids were extracted from each sample according to a modified Bligh and Dyer protocol at 4 °C. Briefly, 1 mL of MeOH was added to each sample (380 µL), supplemented with 4 µL of a deuterated internal standard (SPLASH lipidomix, Avanti Polar), and the mixture was vortexed for 30 sec. Then, 500 µL of cold CHCl_3_ was added, and the sample was mixed for 30 sec. Subsequently, 500 µL of H_2_0 and 500 µL of additional CHCl_3_ were added, and the mixture was vortexed again for 30 sec. Samples were centrifuged at 1690 *x g* for 10 min at 4 °C to induce phase separation. The organic phase (about 1 mL), containing the lipids, was recovered, dried under a nitrogen stream, and stored at -20°C under argon.

Prior to analysis, extracts were resuspended in 25 µL of MeOH: CHCl3 (1:1, v/v). A 3 µL aliquot of each lipid extract was injected on an Accucore C18 reverse-phase liquid chromatography column (150 × 2.1, 2.5µm Thermo Fisher Scientific) connected to a UPLC system (Ultimate 3000, Thermo Fisher Scientific). The column was equilibrated with ACN/H_2_O (1:1, v/v) solution containing 10 mM ammonium formate and 0.1% formic acid (eluent A). Lipid were eluted at a flow rate of 0.4 mL/min with a isopropanol/ACN/H2O (88:10:2, v/v) mix containing 2 mM ammonium formate and 0.02% of formic acid (eluant B) using the following gradient : 0 to 35% B from 0 to 4 min, 35 to 60% B from 4 to 8 min, 60 to 70% B from 8 to 16 min, 70 to 85% B from 16 to 25 min, 85 to 97 from 25 to 25.1 min, 97 to 100 %B from 25.1 to 31 min. The percentage of B decreased to 35% in 4 min. Lipids were detected using a Q-Exactive Mass Spectrometer (Thermo Fisher Scientific) equipped with a heated electrospray ionization (HESI) probe. The spectrometer was controlled by Xcalibur 4.3 and operated in y positive-ion mode.

For each sample, two LC-MS injections were performed: one in Parallel Reaction Monitoring (PRM) mode and one in data-dependent MS^2^ (ddMS^2^) mode. One blank injection was performed between samples. TPRM spectra were acquired at a resolution of 35,000 with an automatic gain control (AGC) target of 2×10^5^ and a maximum injection time (max IT) of 100 ms. MS-MS was triggered with an inclusion list containing the precursor ion masses presented in Table S2 with an isolation window of 1.0 m/zIn ddMS^2^ mode, full MS spectra were acquired at a resolution of 70,000 (at 200 m/z) in a mass range of 250–1200 m/z with an AGC target of 1×10^6^ and a max IT of 100 ms. The 15 most intense precursor ions were selected, isolated with a window of 1.0 m/z, and fragmented by Higher-energy Collisional Dissociation (HCD) with normalized collision energy (NCE) of 25 and 30 %. MS/MS spectra were acquired at a resolution of 35000 (at m/z 200), with an AGC target of 1×10^5^, and an injection time of 80 ms.

PRM peaks were integrated with Xcalibur 4.3 (Quan) based on the corresponding mass filters and the product ions listed in Table S2. Full MS-ddMS^2^ spectra were processed using LipidSearch 5.0 (Thermo Fisher Scientific). Product ion search mode was used, and identifications were based on the accurate mass of precursor ions and MS^2^ spectral patterns. The parameters for the ion identification and alignment were set up as follows: ion precursor tolerance 5 ppm; ion product tolerance 5 ppm; alignment retention time tolerance 0.4 min; ID quality filters A, B, and C. Grade C lipids were also filtered out based on their ID score.

### Flotation assay

Flotation assays using liposomes were performed as described in ^54^ with minor modifications to measure the association of Sfh3 with membranes. The protein (0.75 µM) was incubated with liposomes (750 µM lipids) of the desired composition, doped with 0.1% NBD-PE, for 1 h at 25 °C under constant shaking (800 rpm). The liposome/protein mixture (final volume 150 µL) was adjusted to 28% (w/w) sucrose by adding 100 µL of a 60% (w/w) sucrose solution in HK buffer and overlaid with 200 µL of HK buffer containing 24% (w/w) sucrose and 50 µL of sucrose-free HK buffer. The sample was centrifuged at 201,600 × *g* (average centrifugal force) in a swinging-bucket rotor (TLS 55, Beckman Coulter) for 70 min. The bottom (250 µL), middle (140 µL), and top (110 µL) fractions were collected. The bottom and top fractions were analyzed by SDS-PAGE after staining with SYPRO® Orange using a FUSION FX fluorescence imaging system.

### Nano differential scanning fluorimetry

The relative thermal stability of the Sfh3:ligand complexes was assessed by nanoDSF using a Prometheus NT.48 instrument (NanoTemper GmbH). To prepare the complexes, liposomes (4 mM), composed only of DOPC or additionally containing 5% PI, 18:1/18:1-PI4P, DHE, lanosterol, or zymosterol, were prepared in 50 mM HEPES, pH 7.4, 220 mM sucrose buffer. After extrusion, a suspension of these liposomes (600 µL) was diluted with HK buffer to a final volume of 2 mL and ultracentrifuged for 20 minutes at 542,230 × *g* and 20 °C, using a fixed-angle rotor (TLA 110, Beckman Coulter), to pellet them. The supernatant was removed quickly and carefully, then the liposomes were resuspended in 600 µL of HK buffer. Next, Sfh3 (15–20 μM) was incubated with these liposomes (with at least a 1:15 protein:ligand ratio) in HK buffer to a final volume of 360 μL for 40 minutes at 30 °C under agitation (1000 rpm). After incubation, the samples were centrifuged at 434,902 × *g* for 20 minutes at 20 °C using an angular rotor (TLA 120.1) to pellet the liposomes. Then, 260 μL of the supernatant containing the empty Sfh3 (prepared with DOPC-only liposomes) or lipid-loaded Sfh3 protein was collected. The concentration of Sfh3 was determined for each sample by spectrophotometry. Finally, the protein samples were rapidly frozen using liquid nitrogen and then stored at –80°C. On the day of the experiment, standard capillaries were filled with 10 µL of empty Sfh3 or Sfh3:ligand complex solutions at a 10 µM protein concentration (*i.e*., 0.4 mg/mL). The unfolding profiles were recorded by monitoring the intrinsic fluorescence of Sfh3 at λ_em_ = 330 nm and λ_em_ = 350 nm during a thermal ramp from 25 to 80 °C at a 1.5°C/min rate. The excitation power was set to 60%. The inflection points (IPs) of the unfolding profiles, which represent the F_350_/F_330_ fluorescence ratio as a function of temperature, were determined using the Nanotemper analysis software. As an IP value corresponds roughly to the melting temperature (Tm) at which half of the protein was unfolded, this allows calculation of a difference in T_m_ (ΔT_m_) between Sfh3 in complex with a given lipid ligand and the empty form of Sfh3 prepared with liposomes composed only of DOPC. This difference is representative of the relative thermal stability of complexes. Values are the mean of three replicates. The unfolding profile first derivatives were calculated by the Nanotemper analysis software and allow a representation that makes it easier to visualize the inflection point (*i.e.,* the minimum value of the curve in our case).

### Molecular docking

Docking studies were performed using the open conformation of Sfh3 (PDB ID 4J7Q, chain A)^20^. The structure was preprocessed by removing co-crystallized ligands and water molecules, adding polar hydrogens, and assigning partial charges using MGLTools (https://ccsb.scripps.edu/mgltools/), which was subsequently converted into .pdbqt format. To prepare 16:0/18:2-PI, 16:0/18:2-PI4P, and 18:1/18:1-PI4P, SMILES were adapted from the closest related available structure from PubChem. Then, for each compound, three-dimensional coordinates were generated with the ETKDGv3 method ^55^ as implemented in RDKit (https://www.rdkit.org, v2026.03.6), and the structures were energy-minimized with the MMFF94 force field^56^. Because the stereochemistry of the inositol ring was not reliably reproduced by the RDKit tool, the inositol moiety was taken from the crystallographic PI4P ligand 2Y5 (PDB 4PH7)^57^ and joined to the acyl chains of the RDKit-generated models to make a proper *myo*-inositol-like structure, which actually has 1 axial and 5 equatorial hydroxyl (-OH) groups in its most stable chair conformation. The bridging oxygen of the inositol was retained when the head group was excised, so that the two fragments were linked by forming the natural phosphodiester bond in RDKit rather than by repositioning atoms. For PI, the 4-phosphate was removed from this head group, leaving a free hydroxyl. The transferred head-group atoms were then held fixed while the remainder of each molecule was energy-minimized, so that all three ligands share the experimentally determined head-group geometry. Docking was performed with AutoDock Vina (version 1.1.2)^58^. The grid box was centered on the lipid-binding pocket at -18.513, 7.828, 4.657 with dimensions 28 × 30 × 34 Å, an exhaustiveness of 32 (which allows extensive conformational sampling for the ligand during the docking), and 20 binding modes retained per ligand. Ligands were prepared with AutoDockTools^59^, with polar hydrogens fixed and rotatable bonds assigned automatically. The top-ranked pose of each ligand was selected for structural analysis and merged with the receptor coordinates. Protein–ligand contacts were analyzed using LigPlot+ v2.3.1^60^. Hydrogen bonds were assigned with a maximum donor-acceptor distance of 3.35 Å and a maximum hydrogen–acceptor distance of 2.70 Å, and hydrophobic contacts were assigned over a range of 2.90 to 3.90 Å^61^. Three-dimensional representations were prepared using the PyMOL Molecular Graphics System, Version v3.1.8 Schrödinger, LLC.

### Yeast manipulation and growth assays

*Saccharomyces cerevisiae* strains were grown in rich medium (YPD) containing 2% (w/v) glucose or in synthetic dextrose (SD) medium containing 2% (w/v) glucose and appropriate amino acid drop-out mix (MP Biomedicals). Strains were transformed by the standard lithium acetate/polyethylene glycol procedure. All yeast strains were derived from the BY4741 background (*MATa his3*Δ1 *leu2*Δ0 *met15*Δ0 *ura3*Δ0), with the *sfh3*Δ strain obtained from the Euroscarf deletion collection (*MATa his3*Δ1 *leu2*Δ0 *met15*Δ0 *ura3*Δ0 *sfh3*Δ::KanMX). Plasmids and associated yeast strains are listed in Table S3. The Gibson Assembly Hifi Kit (Ozyme) was used to generate the plasmid pSfh3, and the corresponding mScarlet-tagged plasmids, with the mScarlet sequence amplified from Addgene 179067 (a gift from Vincent Martin). A site-directed mutagenesis kit (Agilent) was used for generating point mutations in Sfh3-mutant-expressing plasmids. Briefly, plasmids were amplified using overlapping oligonucleotides containing the desired mutations, designed with the most frequent codon usage. All constructs were verified by DNA sequencing. These plasmids were then used as templates to generate the PCR fragments used to replace the *KanMX* cassette at the *SFH3* locus. To generate WT and mutant Sfh3-mScarlet-tagged strains in *sfh3*Δ background, we employed a marker-less CRISPR-Cas9 genomic editing system targeting the *KanMX* antibiotic-resistance cassette^62^, inserted at the *SFH3* locus. Donor PCR fragments containing the corresponding WT or mutant *SFH3*-mScarlet sequence were amplified from the respective plasmids and used for CRISPR/Cas9-mediated replacement of the *KanMX* cassette, while retaining the endogenous promoter and 3′-UTR regions. Kanamycin-sensitive and mScarlet-tagged colonies were verified by PCR and DNA sequencing. Sfh3^L158Y^, Sfh3^E235A^, and Sfh3^T264W^ mScarlet-tagged strains were subsequently backcrossed to the parental strain BY4742 (*MATα his3*Δ1 *leu2*Δ0 *lys2*Δ0 *ura3*Δ0). For drop-test assays, miconazole (Sigma, PHR1618) was prepared as a 40 mg/mL stock solution in DMSO, aliquoted as 1000x single-use stocks, and stored at -80 °C for up to 6 months. The drug was directly added to the YPD agar plates at 55 °C. Yeasts were grown overnight in YPD at 30 °C, then cells were adjusted to the lowest OD_600_, serially diluted, and 3 µL were spotted onto the appropriate agar media. Plates were incubated for 2–6 days at 37 °C in the dark to prevent the degradation of the light-sensitive drug.

### Statistical analysis

Statistical analyses were performed using Prism (GraphPad). p values < 0.05, < 0.01, and < 0.001 are identified with 1, 2, and 3 asterisks, respectively. Ns: p ≥ 0.05. The number of replicates (n) used for calculating statistics is specified in the figure legends.

## Acknowledgments

AH and this work were supported by ANR-21-CE13-0014 and ANR-23-CE44-0026 grants from the Agence Nationale de la Recherche.

## Data Availability

Raw data are available from the corresponding author(s) upon reasonable request.

## Supplementary Figures

**Figure S1.**
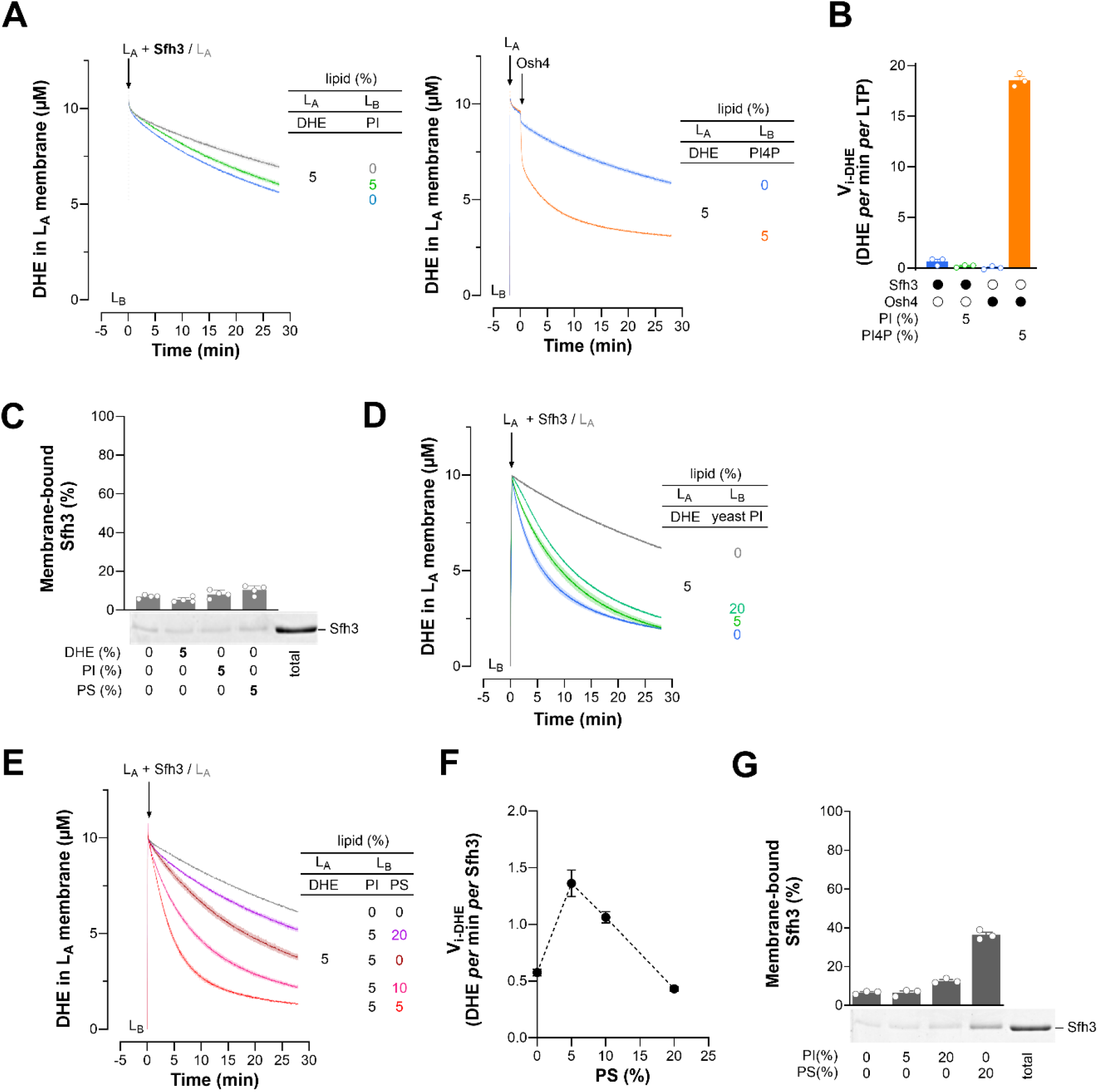
The lipid composition of membranes influences the DHE transfer activity of Sfh3. **(A)** Comparative analysis of the respective influence of PI and PI4P on Sfh3- and Osh4-mediated DHE transfer. Experiments were performed as in Fig. 1D, but with L_A_ and L_B_ liposomes at the same lipid concentration (200 µM). Each trace corresponds to the mean ± s.e.m. of several kinetic traces (n = 3). **(B)** Initial DHE-transfer velocities determined from (A). Data are shown as the mean ± s.e.m. with individual data points (n = 3). **(C)** Flotation assay. Sfh3 (0.75 µM) was incubated in HK buffer at 30 °C for 1 h with agitation in the presence of liposomes (750 µM lipids) composed solely of DOPC or containing 5% PI, 5% PS, or 5% DHE. After centrifugation, liposomes were recovered at the top of sucrose cushions and analyzed by SDS-PAGE. The amount of membrane-bound Sfh3 was quantified relative to lane 5 (100% total) using the SYPRO® Orange signal. Data are presented as mean ± s.e.m. (n = 3), with individual data points. **(D)** Impact of yeast PI levels on Sfh3-mediated DHE transfer. Experiments were performed as in (A), using L_B_ liposomes composed solely of DOPC or containing 5 or 20% yeast PI. Each trace represents the mean ± s.e.m. of several kinetic traces (n = 3–5). **(E)** Influence of the membrane surface charge on Sfh3-mediated DHE transfer. Measurements were performed as in Fig. 1 using L_B_ liposomes composed of DOPC and containing 5% PI, with increasing PS amount (from 0 to 20%) at the expense of DOPC. Control kinetics were recorded without Sfh3, using L_B_ liposomes composed solely of DOPC. Mean ± s.e.m. (with Sfh3, n = 3–4; without Sfh3, n = 2). **(F)** Initial DHE transfer velocity as a function of the percentage of PS in L_B_ liposomes inferred from kinetic data shown in (e). **(G)** Flotation assay. Sfh3 was incubated with liposomes composed of DOPC and containing 5 or 20% PI or 20% PS. Data are presented as mean ± s.e.m. (n = 3), with individual data points.

**Figure S2.**
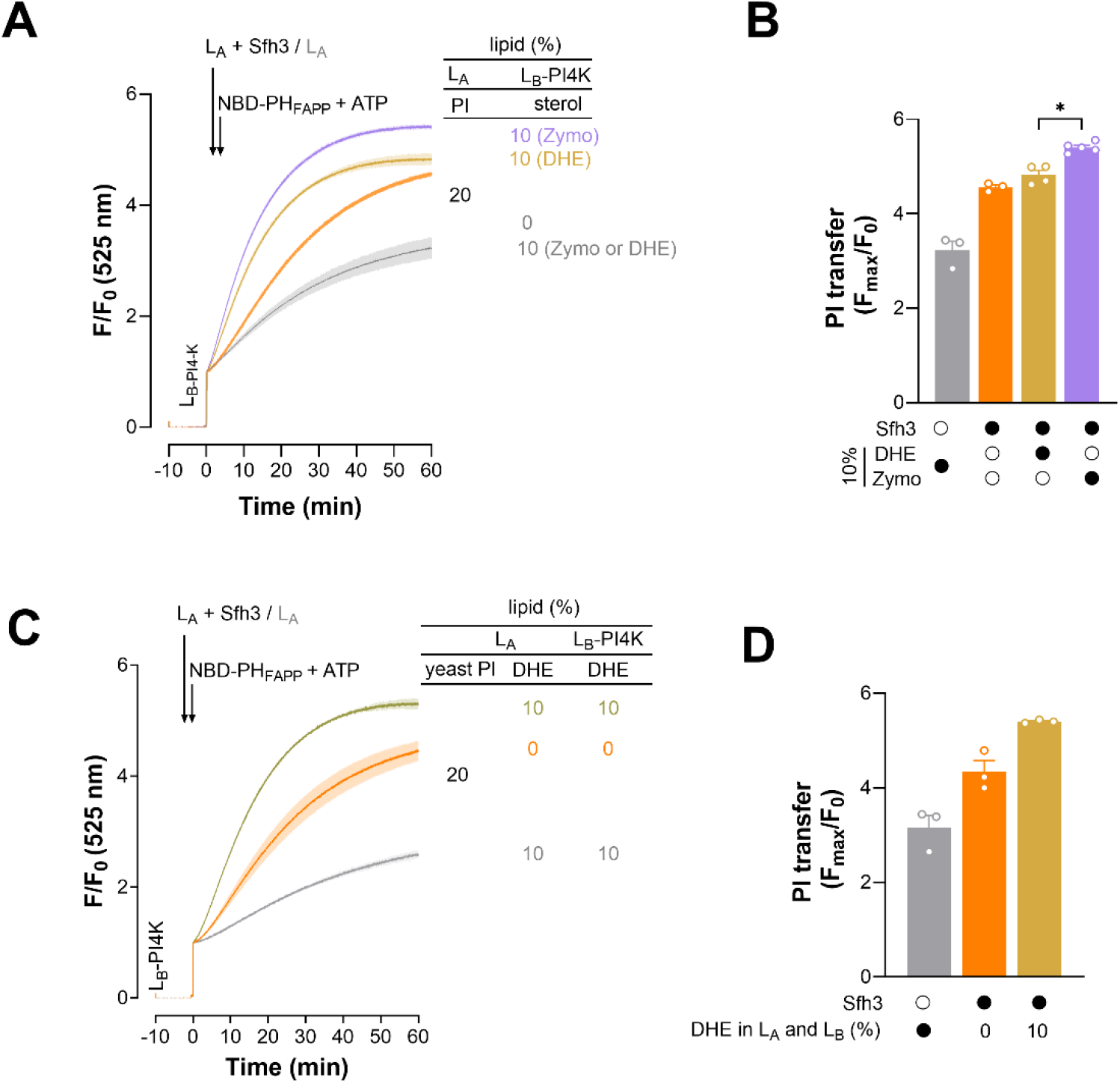
Additional PI transfer assays. **(A)** L_A_ liposomes composed of DOPC/PI/Rhod-PE (78:20:2) were incubated with or without Sfh3 for 1 h at 30 °C, then added to L_B_-PI4K liposomes composed of DOPC/MPB-PE (97:3), DOPC/DHE/MPB-PE (87:10:3), or DOPC/zymosterol/MPB-PE (87:10:3) and coated with PI 4-kinase (L/P = 400). Final concentrations: Sfh3, 200 nM; L_A_ and L_B_ liposomes, 200 µM lipids each. After 30 sec, NBD-PH_FAPP_ (250 nM) and ATP (100 µM) were co-injected. Fluorescence was measured at 525 nm (λ_ex_ = 460 nm) at 30 °C with stirring and normalized to the fluorescence F_0_ measured immediately after the last injection. Each trace is the mean ± s.e.m. of 3–4 kinetic traces. **(B)** PI transfer rates extracted from kinetics shown in (A) as a function of the sterol content of L_B_-PI4K liposomes. Data are represented as mean ± s.e.m. (n = 3–4) with individual data points. Unpaired Mann–Whitney U test (\**p* < 0.05). **(C)** Experiments were performed as in (A) with L_A_ liposomes containing 10% yeast PI instead of soy PI and doped or not with 10% DHE, and L_B_-PI4K liposomes containing or not 10% DHE. A control experiment was performed without Sfh3. Each trace is the mean ± s.e.m. of 3 kinetic traces. **(D)** PI transfer rates extracted from kinetics shown in (D) as a function of the sterol content of L_B_-PI4K liposomes. Data are represented as mean ± s.e.m. (n = 3) with individual data points.

**Figure S3.**
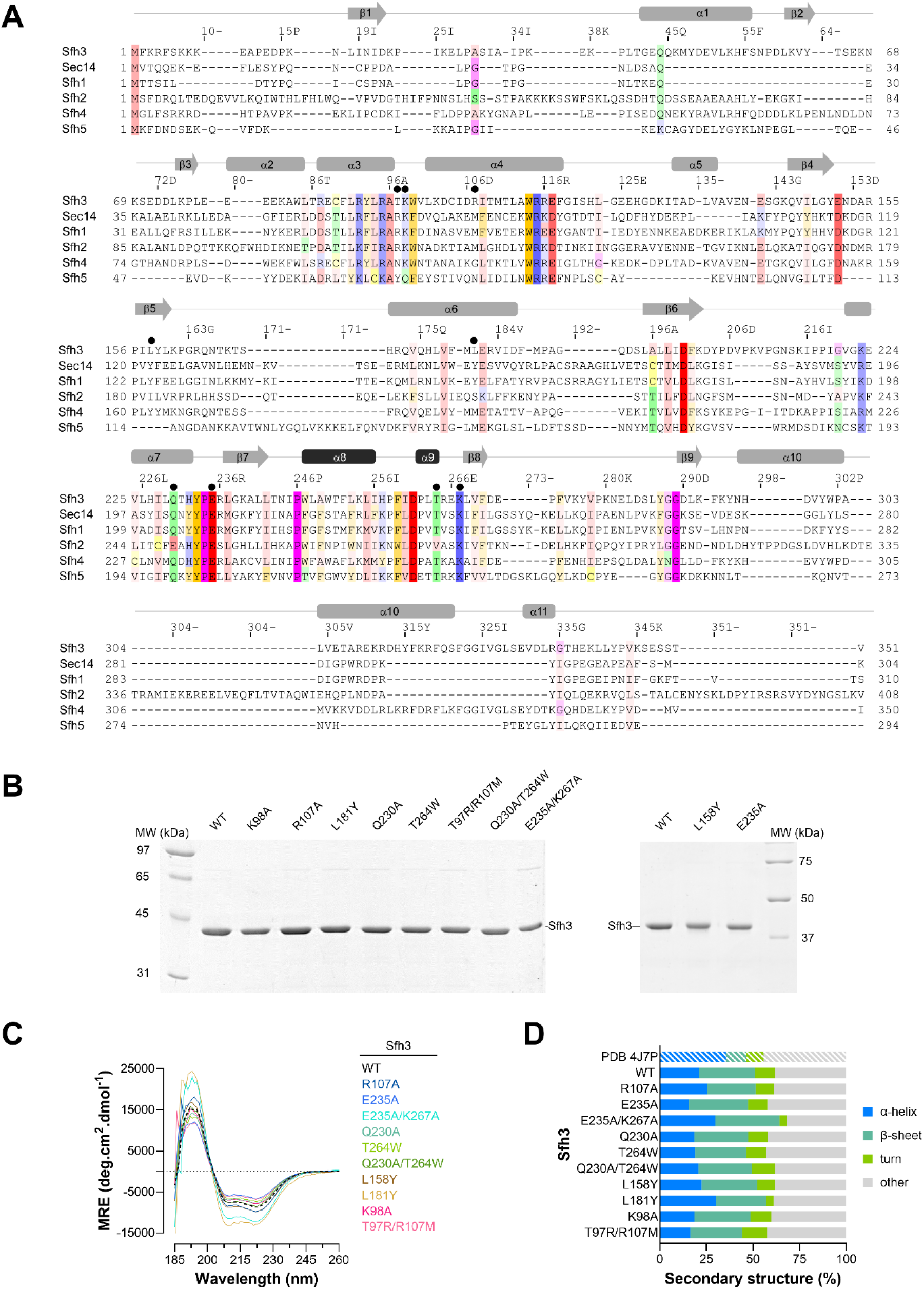
Biochemical and structural analyses of Sfh3 WT and mutants. **(A)** Sequence alignment between Sfh3 (UniProt: P53860), Sec14 (P24280), and the other yeast Sfh proteins (P33324, Q06705, P53844, A6ZQI5). The alignment was performed using T-Coffee. Residues are colored using the Zappo color scheme (pink: aliphatic; orange: aromatic; purple: glycine or proline; yellow: cysteine; green: hydrophilic; red: negatively charged; blue: positively charged) for residues conserved between proteins with a conservation percentage above 40%. The positions of the α-helices and β-sheets along the Sfh3 sequence are shown, with the gating helices α8 and α9 highlighted in black. The positions of the Sfh3 residues that were mutated in our structure-function analysis are indicated with black dots. **(B)** SDS-PAGE analysis of purified Sfh3 WT and mutants. **(C)** Far-UV CD spectrum of purified Sfh3 and mutants (concentration range: 5–14 µM) in 20 mM Tris, pH 7.4, 120 mM NaF buffer at room temperature. MRE, mean residue ellipticity. **(D)** Percentages of secondary structure, derived from the analysis of each spectrum, are provided, along with values derived from the crystal structure (PDB ID: 4J7P) using the BeStSel algorithm.

**Figure S4.**
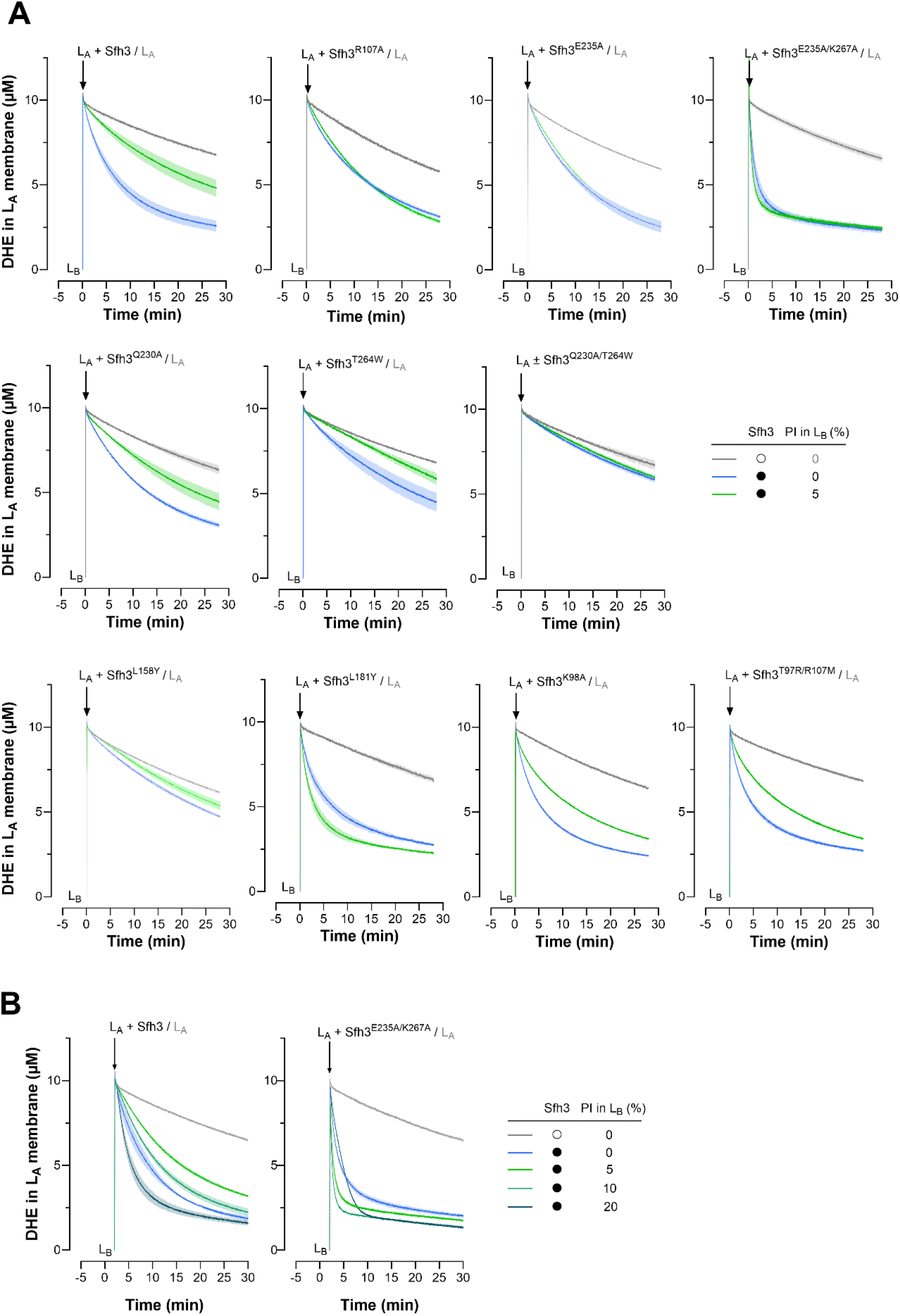
DHE transfer activity of Sfh3 WT and mutants. **(A)** DHE transfer assay. Sfh3 WT or mutants were incubated with L_A_ liposomes composed of DOPC/DHE/DNS-PE (92.5:5:2.5) for 1 h at 30°C. The mixture was then added to L_B_ liposomes containing only DOPC or DOPC plus 5% PI. Fluorescence was measured at 525 nm (λ_ex_ = 310 nm) at 30 °C with constant stirring. The signal was normalized to the DHE amount in the L_A_ membrane, expressed as a concentration. The final concentrations of LTP, L_A,_ and L_B_ liposomes are 1.5, 200, and 600 µM, respectively. Control experiments were performed without LTP, using L_B_ liposomes composed solely of DOPC. Each trace is the mean ± s.e.m. of 3–4 kinetic traces. **(B)** Measurements were performed as in (A) by incubating Sfh3 WT or Sfh3^E235A/K267A^ with L_A_ liposomes composed of DOPC/DHE/DNS-PE (92.5:5:2.5) and then adding the mixture to L_B_ liposomes containing increasing PI amounts (0 to 20%). Each trace is the mean ± s.e.m. of 3–4 kinetic traces.

**Figure S5.**
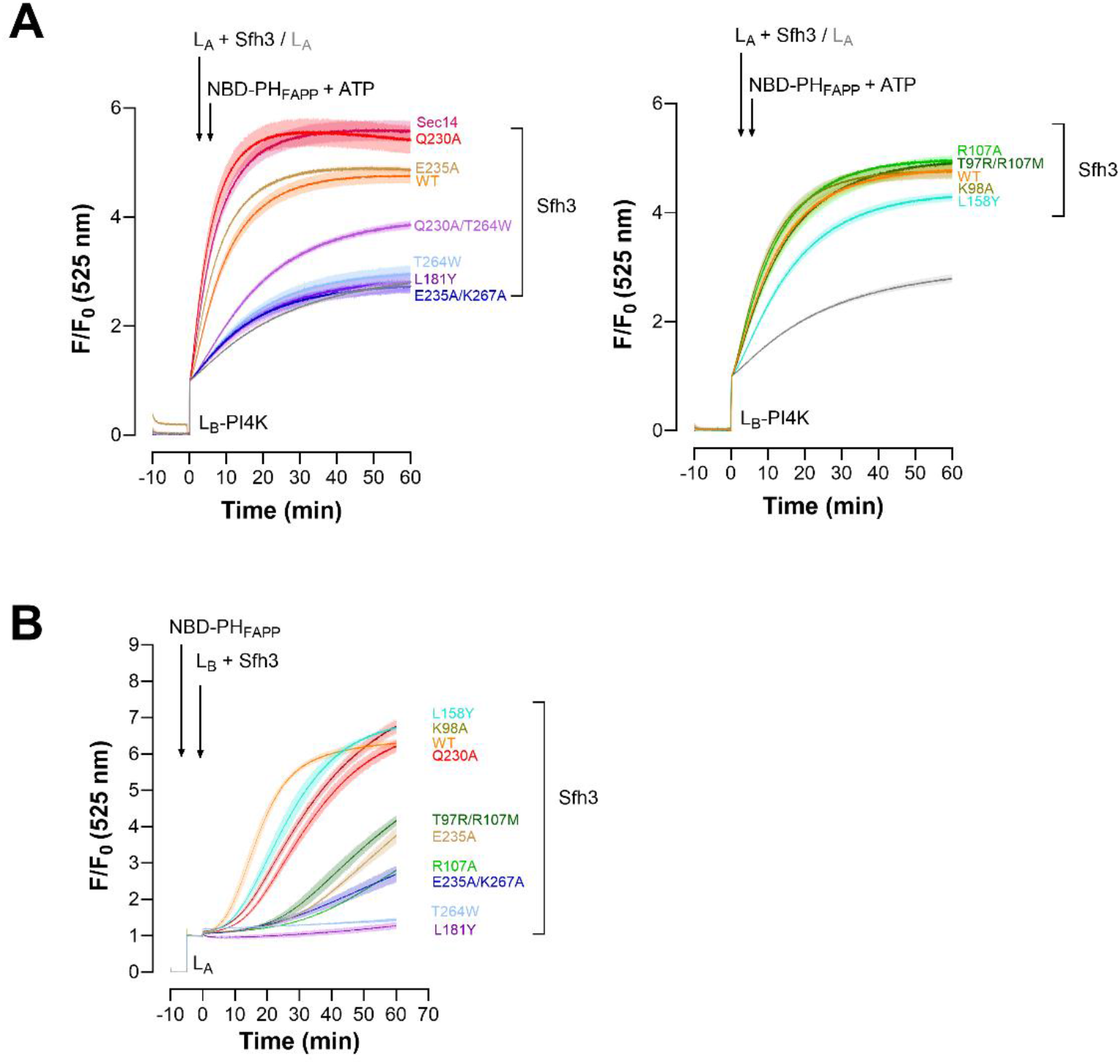
PI and PI4P transfer activities of Sfh3 WT and mutants. **(A)** PI-transfer assay. L_A_ liposomes composed of DOPC/PI/Rhod-PE (78:20:2) were incubated with or without Sec14, Sfh3 WT, or its mutated forms for 1 h at 30 °C and then added to L_B_-PI4K liposomes, composed of DOPC/lanosterol/MPB-PE (87:10:3) and coated with PI 4-kinase (L/P = 400). Final concentration: Sfh3 or Sec14, 200 nM; L_A_ and L_B_ liposomes, 200 µM lipids each. After 30 sec, NBD-PH_FAPP_ (250 nM) and ATP (100 µM) were co-injected. Control kinetics without LTP were recorded. Fluorescence was measured at 525 nm (λ_ex_ = 460 nm) at 30 °C with stirring and normalized to the fluorescence F_0_ measured immediately after the last injection. Each trace is the mean ± s.e.m. of 3–9 kinetics. The curves are shown in two panels for clarity. **(B)** PI4P transfer assays. Sfh3 or each Sfh3 mutant (200 nM) was pre-incubated with L_B_ liposomes composed of DOPC; the mixture was then added to L_A_ liposomes containing 5% PI4P and 2% Rhod-PE, pre-mixed with NBD-PH_FAPP_ (250 nm). The concentration of L_A_ and L_B_ liposomes is 200 µM lipids each. Each trace is the mean ± s.e.m. of 3–5 kinetics.

**Figure S6.**
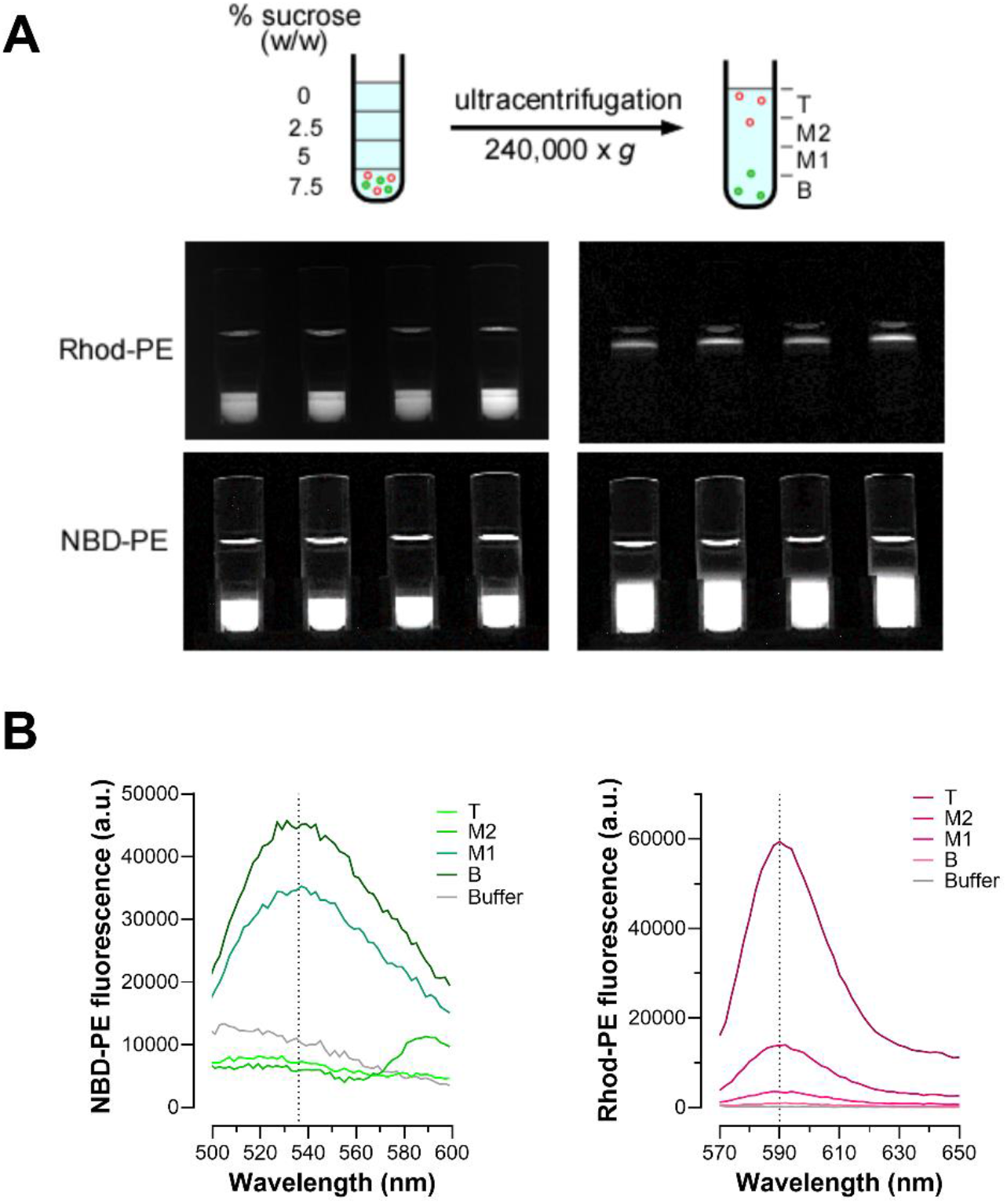
LC-MS/MS-based lipid transfer assays. **(A)** Liposome separation by differential flotation assay. “Heavy” sucrose-loaded liposomes (L_A_, in green) made of DOPC, doped with NBD-PE, containing or not lipid ligand, were pre-mixed or not with Sfh3 and then added to “light” sucrose-free liposomes, composed of DOPC, and including Rhod-PE and 0 or 5% lipid ligand (L_B_, in red) to initiate the transfer experiment. The sample (125 µL) was mixed for 20 minutes at 30 °C under constant agitation. Then, it was transferred to a polycarbonate tube, adjusted to 7.5% sucrose by mixing 75 µL of a 20% (w/w) sucrose HK buffer, then overlaid with three 200 µL layers of 5%, 2.5%, and 0% sucrose HK buffer, respectively. The suspension was centrifuged at 240,000 × *g* for 1 hour, and the bottom (B, 200 µL), middle 1 (M1, 200 µL), middle 2 (M2, 200 µL), and top (T, 200 µL) fractions were collected. Each tube was imaged before and after ultracentrifugation to confirm that the heavy and light liposomes are, respectively, at the bottom and top of the sucrose gradient and are therefore well separated. **(B)** The relative amounts of L_A_ and L_B_ liposomes in B, M1, M2, and T fractions were precisely quantified by mixing 10 µL of each fraction with 90 µL of HK buffer and measuring the maximal fluorescence of NBD-PE and Rhod-PE. The NBD-PE signal was recorded from 495 to 600 nm (maximum at 536 nm) upon excitation at λ_ex_ = 460 nm, whereas the Rhod-PE signal was measured from 570 to 650 nm (maximum at 590 nm) upon excitation at λ_ex_ = 560 nm.

**Table S1.** Protein–ligand interactions in open-state Sfh3 complexes.

| Structure and model | Hydrogen-bonded residues (Å) | Hydrophobic contacts |
| --- | --- | --- |
| Sfh3 :PI (4J7Q, chain A) | R107-Ins 4OH (2.74), E235-Ins 2OH (3.00), R236-Phos1 (2.79), L237-Phos1 (2.85) | A96, Q165, Q173, L177, L181, L197, F201, V225, L229, Y233, P234, L242, A249, K267 |
| Sfh3:16:0/18:2–PI | A96-Ins 4OH (3.01), Q230-Ins 5OH (2.80), Q230-Ins 6OH (2.95), E235-Ins 1OH (2.85), **E235-Ins 2OH (3.07 ,3.23), R236-Phos1 (3.07), L237-Phos1 (2.97)** | K144, Q145, L158, L160, P162, Q165, L177, L181, L195, L197, I199, L229, Y233, P234, L268 |
| Sfh3:16:0/18:2–PI4P | A96-Ins 3OH (3.20), R107-Phos4 (3.07), Q230-Ins 5OH (3.17), E235-Ins 1OH (2.96), **E235-Ins 2OH (3.31), L237-Phos1 (3.27)** | L160, P162, L177, L181, L195, L197, I199, F201, L229, Y233, P234, R236, A240, L242 |
| Sfh3:18:1/18:1–PI4P | A96-Ins 6OH (3.02), R107-Phos1 (3.07, 3.18), **E235-2OH (2.72)**, E235-*sn*2 acyl chain, L237-*sn*2 acyl chain (3.13), R236-*sn*2 acyl chain (2.93) | L160, P162, L177, L181, L195, L197, I199, L229, Q230, Y233, P234, A240, T264, K267, L268 |

Hydrogen-bond donor–acceptor distances are given in Ångströms. Contacts were assigned with identical criteria for all four structures. Residues in bold are those that are found to be in contact with PI in the crystal structure of the Sfh3:PI complex. Ins : inositol ; Phos : phosphate group; OH: hydroxyl function.

**Table S2.** LC-MS/MS detection of main analytes.

| Name | Molecular weight (g/mol) | Retention time (min) | Adduct type | Precursor ion mass (m/z) | Normalized collision energy (%) | Product ion mass (m/z) |
| --- | --- | --- | --- | --- | --- | --- |
| **DHE** | 394.632 | 8.2 | [M+H-H<sub>2</sub>O]<sup>+</sup> | 377.320 | 35 | 157.101<br>69.070 |
| **Lanosterol** | 426.72 | 10.2 | [M+H-H<sub>2</sub>O]<sup>+</sup> | 409.383 | 25 | 109.101<br>191.180 |
| **Zymosterol** | 384.637 | 9.1 | [M+H-H<sub>2</sub>O]<sup>+</sup> | 367.336 | 35 | 95.086<br>81.070 |
| **Cholesterol-D7** | 393.697 | 10.4 | [M+H-H<sub>2</sub>O]<sup>+</sup> | 376.395 | 35 | 243.210<br>161.132 |
| **NBD-PE** | 956.24 | 9.3 | [M+H]<sup>+</sup> | 924.581 | 25 | 603.530<br>339.289 |
| **Rhod-PE** | 1301.71 | 8.8 | [M+H-H<sub>2</sub>O]<sup>+</sup> | 1284.692 | 25 | 682.164<br>602.198 |
| **PE-D7** | 711.013 | 10.6 | [M+H]<sup>+</sup> | 711.566 | 35 | 570.547 |

**Table S3.**
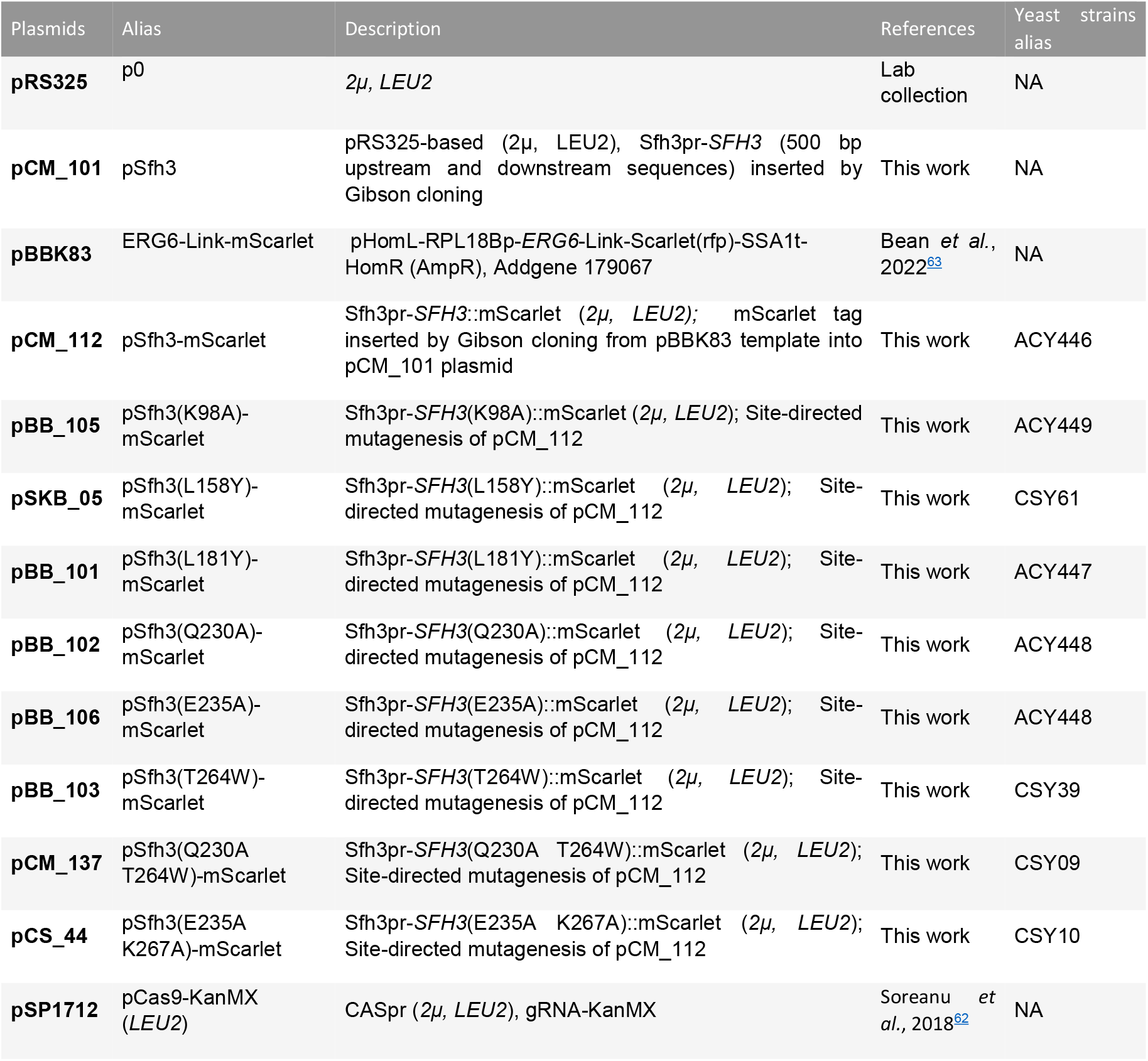
Plasmids and yeast strains used in this study. Note that all plasmids carry ampicillin antibiotic resistance.

| Plasmids | Alias | Description | References | Yeast strains |
| --- | --- | --- | --- | --- |
| **pRS325** | p0 | *2μ, LEU2* | Lab collection | NA |
| **pCM_101** | pSfh3 | pRS325-based (*2μ, LEU2*), Sfh3pr-*SFH3* (500 bp upstream and downstream sequences) inserted by Gibson cloning | This work | NA |
| **pBBK83** | ERG6-Link-mScarlet | pHomL-RPL18Bp-*ERG6*-Link-Scarlet(rfp)-SSA1t-HomR (AmpR), Addgene 179067 | Bean *et al.*, 2022<sup>63</sup> | NA |
| **pCM_112** | pSfh3-mScarlet | Sfh3pr-*SFH3*::mScarlet (*2μ, LEU2*); mScarlet tag inserted by Gibson cloning from pBBK83 template into pCM_101 plasmid | This work | ACY446 |
| **pBB_105** | pSfh3(K98A)-mScarlet | Sfh3pr-*SFH3*(K98A)::mScarlet (*2μ, LEU2*); Site-directed mutagenesis of pCM_112 | This work | ACY449 |
| **pSKB_05** | pSfh3(L158Y)-mScarlet | Sfh3pr-*SFH3*(L158Y)::mScarlet (*2μ, LEU2*); Site-directed mutagenesis of pCM_112 | This work | CSY61 |
| **pBB_101** | pSfh3(L181Y)-mScarlet | Sfh3pr-*SFH3*(L181Y)::mScarlet (*2μ, LEU2*); Site-directed mutagenesis of pCM_112 | This work | ACY447 |
| **pBB_102** | pSfh3(Q230A)-mScarlet | Sfh3pr-*SFH3*(Q230A)::mScarlet (*2μ, LEU2*); Site-directed mutagenesis of pCM_112 | This work | ACY448 |
| **pBB_106** | pSfh3(E235A)-mScarlet | Sfh3pr-*SFH3*(E235A)::mScarlet (*2μ, LEU2*); Site-directed mutagenesis of pCM_112 | This work | ACY448 |
| **pBB_103** | pSfh3(T264W)-mScarlet | Sfh3pr-*SFH3*(T264W)::mScarlet (*2μ, LEU2*); Site-directed mutagenesis of pCM_112 | This work | CSY39 |
| **pCM_137** | pSfh3(Q230A T264W)-mScarlet | Sfh3pr-*SFH3*(Q230A T264W)::mScarlet (*2μ, LEU2*); Site-directed mutagenesis of pCM_112 | This work | CSY09 |
| **pCS_44** | pSfh3(E235A K267A)-mScarlet | Sfh3pr-*SFH3*(E235A K267A)::mScarlet (*2μ, LEU2*); Site-directed mutagenesis of pCM_112 | This work | CSY10 |
| **pSP1712** | pCas9-KanMX (*LEU2*) | CASpr (*2μ, LEU2*), gRNA-KanMX | Soreanu *et al.*, 2018<sup>62</sup> | NA |

